# The *Escherichia coli* replication initiator DnaA is titrated on the chromosome

**DOI:** 10.1101/2024.10.07.617004

**Authors:** Lorenzo Olivi, Stephan Köstlbacher, Mees Langendoen, Nico J. Claassens, Thijs J.G. Ettema, John van der Oost, Pieter Rein ten Wolde, Johannes Hohlbein, Raymond H. J. Staals

**Affiliations:** Laboratory of Microbiology, Wageningen University & Research, Wageningen, the Netherlands; Institute AMOLF, Amsterdam, the Netherlands; Laboratory of Biophysics, Wageningen University & Research, Wageningen, the Netherlands; Microspectroscopy Research Facility, Wageningen University & Research, Wageningen, the Netherlands

## Abstract

DNA replication initiation is orchestrated in many prokaryotes by the replication initiator DnaA. Two models for regulation of DnaA activity in *Escherichia coli* have been proposed: the switch between an active and inactive form of DnaA, and the titration of DnaA on the *E. coli* chromosome. Although proposed decades ago, experimental evidence of a titration-based control mechanism is still lacking. Here, we first identified a conserved high-density region of binding motifs near the origin of replication, an advantageous trait for titration of DnaA. We then investigated the mobility of DnaA by single-particle tracking microscopy in wild-type and deletion mutants *E. coli* strains, while monitoring cellular size and DNA content. Our results indicate that the chromosome of *E. coli* controls the free amount of DnaA in a growth rate-dependent fashion. Finally, we provide insights on the relevance of DnaA titration in stabilising DNA replication by preventing re-initiation events during slow growth.

## Introduction

The correct coordination of DNA replication with cellular growth is crucial for any living organism^1^. Remarkably, the orchestration of these processes allows *Escherichia coli* to divide faster than the time it takes to replicate its chromosome^2–5^. This surprising ability was first explained in 1968 by a model stating that *E. coli* initiates one or more rounds of replication before the initial replicative event is completed^2^. Subsequently, an extension of the model introduced the concept of volume of initiation (*V*\*), a specific cellular size per origin of replication, the chromosomal locus at which the DNA replication process is initiated^6^. Since then, the coupling of DNA replication and cellular growth has been experimentally confirmed^4,5^ and the initiation of DNA replication is now considered to be associated with the accumulation of a constant amount of cellular volume per origin^7–9^. The volume of initiation is one of the most tightly controlled parameters in the *E. coli* cell cycle^5,10,11^.

At the molecular level, *E. coli* initiates DNA replication by unwinding the origin of replication (*oriC*) through the binding of the replication initiator protein DnaA^12^. The regulation and biochemistry of DnaA has been extensively studied^12,13^, yet how *E. coli* achieves its characteristic DNA replication behaviour is still a matter of debate^1,12,14–16^. At the expression level, balanced biosynthesis of the *dnaA* gene generates stable intracellular concentrations of DnaA across the cell cycle^17,18^. Moreover, the expression of *dnaA* is negatively auto-regulated, with DnaA repressing its own promoter *DnaAp2*^19^. This type of regulation, together with balanced biosynthesis, may explain that concentrations of DnaA vary of less than 50% over a tenfold change in growth rate^11^. DnaA binds DNA at 9 bp-long sequences, termed DnaA boxes. Both high-affinity and low-affinity boxes exist, with the first class matching a consensus sequence (TTWTNCACA)^20^ and the latter carrying multiple variations^21,22^. Two classes of models have been proposed to clarify how the DnaA-mediated unwinding of *oriC* is regulated. First, the initiator titration model^23,24^ dictates that DnaA is sequestered by hundreds of high-affinity DnaA boxes present on the chromosome of *E. coli*^15^ (Figure 1, top). Only after these boxes have been saturated, DnaA can bind the low-affinity boxes present at *oriC* and unwind it, thereby initiating DNA replication. When structural and biochemical characterisation of DnaA advanced, a second model emerged. This switch model (Figure 1, bottom) is based on the finding that DnaA switches between an ATP-bound state competent in origin unwinding, and an inactive, ADP-bound state^12^. The interconversion between states is regulated by several mechanisms and when the levels of DnaA-ATP reach a threshold replication is initiated. Newly synthesised DnaA proteins predominantly bind to ATP due to its higher abundance in the cell compared to ADP^25^. Further, non-coding sequences called DnaA-Reactivating Sequences 1 and 2 (*DARS1* and *DARS2*) are involved in recruiting DnaA to nine particular DnaA boxes, where ADP is replaced with ATP^26^. At the same time, a similar re-activating mechanism has been proposed for interactions with acid phospholipids *in vitro*^27^, which however is still lacking confirmation *in vivo*. Conversely, the non-coding sequence *datA* recruits DnaA with five other DnaA boxes, where *datA-*mediated DnaA-ATP hydrolysis (DDAH) occurs^28^. Finally, the Hda protein associates with the β-clamp of DNA polymerase III during replication to stimulate hydrolysis of DnaA-bound ATP in a process termed regulatory inactivation of DnaA (RIDA)^29^. Moreover, both DnaA-ATP and DnaA-ADP have been shown to be competent in binding high-affinity boxes, whereas only DnaA-ATP can bind to low-affinity boxes^21,22,30^.

**Figure 1.**
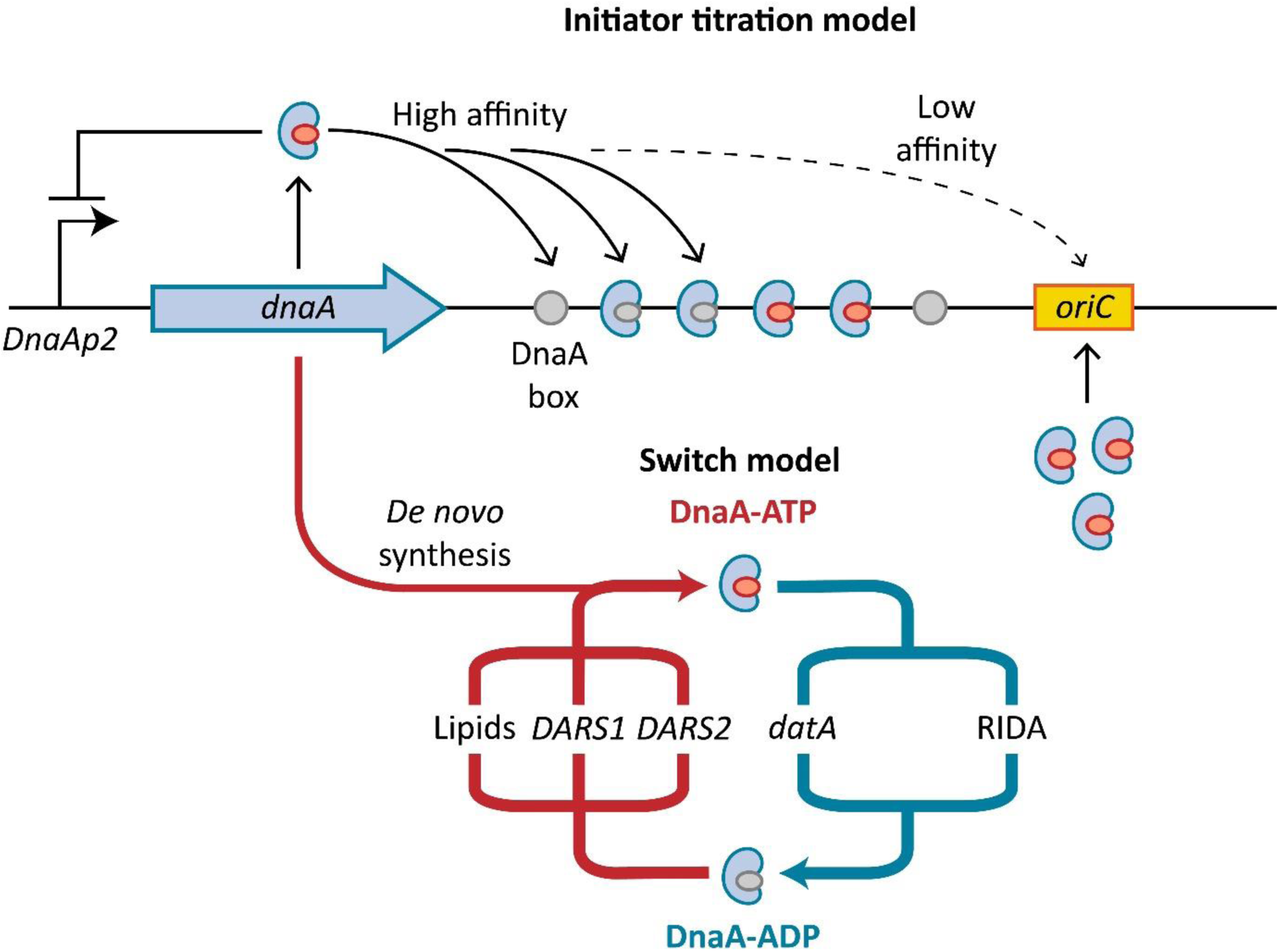
The two models attempting to explain the control of DNA replication in *E. coli*. DnaA negatively auto-regulates its own expression (left). Newly synthesised DnaA is bound by ATP. In the *initiator titration model* (top), the initiator protein is sequestered by high-affinity DnaA boxes throughout the chromosome. Only when all boxes are saturated, DnaA can bind to low-affinity boxes present on *oriC*, initiating DNA replication. In the *switch model* (bottom), DnaA is present in either an ATP-bound, active form, or an ADP-bound, inactive form. Several mechanisms are involved in the switch between the two forms. Once DnaA-ATP accumulates over a specific threshold, it can bind and unwind *oriC*.

*E. coli* strains carrying deletions of the switch control components *datA*, *DARS1, DARS2* and *hda,* either individually or combined, exhibit defects in initiation, but are still viable^18,31,32^. Therefore, other forms of control must be present. The viability of the quadruple mutant was attributed to the intrinsic ATPase ability of DnaA, while postulating that initiator titration can provide an additional stabilising effect^18^. Previous *in silico* work also suggested that concerted DnaA interconversion and its titration on the chromosome can enhance the stability of *E. coli* cell cycles in all growth conditions^33,34^, with a larger effect at lower growth rates^35^.

The titration of DnaA on the *E. coli* chromosome has been first hypothesised more than 40 years ago^23^ and has been considered as being present in many recent modelling^33–35^ and experimental studies^18,32^. Yet, there is still no experimental evidence that *E. coli* can control the free concentration of DnaA via titration. To address this gap in knowledge, we first analysed the configuration of *E. coli* genome and showed that DnaA boxes are distributed in a way that would allow titration. Specifically, the DnaA binding motifs are accumulated towards the origin of replication, a trait that is conserved within many *E. coli* strains as well as in *Salmonella enterica*. On an experimental level, revealing the presence of DnaA titration requires assessing the binding state of this protein in live cells. Previous biochemical assays already probed the interactions of DnaA with DNA^20,36,37^, but the required lysis step inevitably caused all cell cycle-dependent variation to be lost in favour of population-level measurements. Here, we overcame this limitation by employing single-particle tracking photoactivatable localisation microscopy (sptPALM)^38^ on DnaA in live *E. coli* cells. We generated fusions of the native DnaA with the photoactivatable fluorescent protein PAmCherry2.1^39^ in the wild-type *E. coli* MG1655 and in mutant strains in which *datA, DARS1* and *DARS2* were deleted either individually or together^40^. Building on our previously established experimental pipeline^39^ (Supplemental Figure 1), we grew all strains in constant cellular density settings, monitored relevant cellular parameters (i.e., cellular area and DNA content) and performed sptPALM. We calculated the overall bound fraction of DnaA, as well as the mobility of the DnaA tracks generated through sptPALM throughout the cell cycle in a variety of different growth conditions. Our data provide experimental evidence that the *E. coli* chromosome is indeed capable of titrating DnaA, keeping its diffusivity low in a growth-rate-dependent fashion. The analysis presented here suggests a role for the initiator titration mechanism in stabilising *E. coli* DNA replication initiation, preventing premature re-initiation events.

## Materials and methods

### Computational identification of chromosomal DnaA box densities

To analyse the density of DnaA boxes across the chromosome of *E. coli,* nucleotide fasta files of several *E. coli* strains genomes^40^, as well as the reference genome of *Salmonella enterica* were downloaded from RefSeq^41^. Identifiers NC_017644.1, NC_017663.1, and NC_020518.1 were removed from RefSeq and therefore excluded from analysis. A custom-made Python script (matchBox.py) was developed to scan circular chromosome and identify exact matches to the DnaA box motifs, either on the forward or reverse strand. The consensus DnaA box sequence (TTWTNCACA) and the sequence of other conserved boxes present in *datA, DARS1* and *DARS2* were included in the list of potential matches. We accounted for the circular nature of *E. coli* and *S. enterica* chromosomes by allowing detection of motifs near the origin and terminus. Matches were identified and assigned a sequence ID, position, orientation and motif. A complete list of matches is available by contacting the authors, together with all the custom-written scripts used in the study. The distribution of DnaA box motifs was then analysed and visualised with R v4.4.1^42^. The relative positions of DnaA box motifs within each chromosome was calculated by scaling to centisomes, i.e. dividing the total length of each genome in 100 equal intervals^43^, starting from the position of *oriC,* determined via the DoriC database^44^.

We then tested whether DnaA boxes are distributed uniformly over the chromosome. We generated the null hypothesis of DnaA boxes being homogenously distributed over the chromosome by repeatedly (*n* = 1000) simulating DnaA box positions over the chromosome from a uniform distribution in R v4.4.1^42^ with the ‘runif’ function. We calculated the number of boxes per position in a sliding window approach by moving over the scaled chromosome in a 10 centisome window, taking 1 centisome steps. We then calculated pseudo *p*-values based on the rank of the real box number per position compared to the simulated values from a uniform distribution with a significance cutoff of *p*-value < 0.05.

Additionally, we tested whether the non-uniform distribution of DnaA boxes derived from the genome compositional background (i.e. GC content), or if these sequences were effectively overrepresented. To this end, we created 1,000 permutations of the consensus DnaA box sequence (TTWTNCACA)^20^ and of other sequences reported as active in DnaA binding across *datA, DARS1* and *DARS2* (HHMTHCWVH)^40^. We searched the chromosome for identical matches to the permutated box sequences and calculated the number of motifs per position in a sliding window approach as for the uniform distribution test. Pseudo *p*-values were calculated and used as before to obtain significance cutoffs.

Circular statistics was then employed to estimate the density of DnaA boxes along the genomes, using the circular v0.5-0 package^45^, and the von Mises kernel to smooth the density of motifs. The density estimates were subsequently scaled to Z-score to indicate the number of standard deviations the value is away from 0, i.e. the mean density of boxes across the chromosome. Extreme density values were extracted with the zoo v1.8-12 package^46^ ‘rollapply’ function, with the largest Z-score assigned as the global maximum for that genome. Data was then visualised with the ggplot2 v3.5.1 package^47^.

### Bacterial strains, plasmids and growth conditions

All bacterial strains and relative genotypes used in this study are described in Supplemental Table 1. *Escherichia coli* DH5α (NEB) was used for general plasmid propagation and standard molecular techniques. The wild-type *Escherichia coli* MG1655 and mutant strains lacking the genomic loci *datA, DARS1* and *DARS2* either individually or in combination^40^ were used to investigate DnaA interactions with the bacterial chromosome.

All *E. coli* strains were routinely grown at 30 °C or 37 °C and 200 rpm in LB liquid medium (10 g/L tryptone, 10 g/L NaCl, 5 g/L yeast extract) for strain propagation. M9 liquid medium (5 g/L KH_2_PO_4_, 0.5 g/L NaCl, 6.78 g/L Na_2_HPO_4_, 1 g/L NH_4_Cl, 4.98 mg/L FeCl_3_, 0.84 mg/L ZnCl_2_, 0.13 mg/L CuCl_2_ · 2H_2_O, 0.1 mg/L CoCl_2_ · 6H_2_O, 0.1 mg/L H_3_BO_3_, 0.016 mg/L MnCl · 4H_2_O, 0.5 g/L MgSO_4_ · 7H_2_O, 11 mg/L CaCl_2_) was used during constant density cultivations or single-particle tracking. Growth conditions to achieve different growth regimes were: M9 supplemented with 4 g/L glucose and 1x RPMI 1640 amino acids at 37 °C for fast growth regime; M9 supplemented with 4 g/L succinate and 1x RPMI 1640 amino acids at 30 °C for intermediate growth regime; and M9 supplemented with 4 g/L acetate at 37 °C for slow growth regime. Media were eventually supplemented with agar (15 g/L) to obtain solid media and with antibiotics for plasmid propagation (50 mg/L kanamycin, 100 mg/L spectinomycin, 100 mg/L ampicillin, 25 mg/L chloramphenicol).

Phosphate-buffered saline solution (PBS) (8 g/L NaCl, 0.2 g/L KCl, 1.42 g/L Na_2_HPO_4_, 0.24 g/L KH_2_PO_4_) was used to wash cells during sample preparation for single-particle tracking experiments.

### Plasmid construction

All constructs used in this study are described in Supplemental Table 2. The pCas and pTarget system^48^ was used to obtain genomic mutants of *E. coli* MG1655. The pTarget_dnaA-PAFP plasmid series were constructed from pTarget^48^. The pTarget_dnaA-PAFP plasmid series expressed a single-guide RNA targeting the chromosomal *dnaA* gene of *E. coli* in its domain II and a repair template flanked by 50 bp-long homology arms. The repair template consisted of the gene encoding one of four photoactivatable or photoconvertible fluorescent proteins (PAFP), replacing bases 259-312 of the native *dnaA* gene^49,50^. The tested PAFP were Dronpa2^51^, mEos4b^52^, mMaple3^53^ and PAmCherry2.1, a M10L^54^ mutant of PAmCherry2^55^. Sequences were codon-optimised using the IDT Codon Optimisation Tool (IDT) and DNA fragments coding for Dronpa2 (BG25452), mEos4b (BG25454) and mMaple3 (BG25455) were chemically synthesised (IDT). The sequence of PAmCherry2.1 was retrieved from the pLbdCas12a-PAmCherry2.1_scrambled plasmid^39^.

For cloning purposes, DNA fragments were amplified by PCR using Q5® High-Fidelity DNA Polymerase (NEB) following manufacturer instructions. Specific oligonucleotides were designed and synthesised (IDT) to introduce proper overhangs for assembly. Assembly was performed using NEBuilder® HiFi DNA Assembly Master Mix (NEB). All oligonucleotides and fragments used for cloning purposes are described in Supplemental Table 3.

### Electro-competent cells preparation and transformation procedure

Pre-cultures of *E. coli* were grown overnight in LB liquid medium at 37 °C and 200 rpm. Cells were made electrocompetent by re-inoculating and culturing them at 37 °C and 200 rpm in LB. Strains carrying either the pCas9^48^, the pCas9_ampR or the pSIJ8^56^ plasmid were instead cultured at 30 °C and 200 rpm in LB, supplemented with L-arabinose (final concentration 10 mM) to induce expression of the λ-Red system. Upon reaching OD_600nm_ of 0.4, the cells were cooled down to 4 °C, washed one time with 1 culture volume of ice-cold Milli-Q water and two times with 0.5 culture volumes of an ice-cold 10% V/V glycerol solution in Milli-Q water. Cells were then suspended in ice-cold 10% glycerol to a final volume of 400 µL for each 100 mL of initial culture volume and dispensed in 40 μL aliquots. All washing steps were performed at 4 °C, by centrifugation for 10 min at 3000 x*g*.

Electroporation was performed in ice-cold 2 mm electroporation cuvettes at 2500 V, 200 Ω and 25μF. Immediately after electroporation, cells were recovered in LB medium. Recovery was performed either at 30 °C, 750 rpm for 2 h for cells harbouring either the pCas9, the pCas9_ampR or the pSIJ8 plasmid or at 37 °C, 750 rpm for 1 h in all other instances. Following recovery, cells were spread on LB solid medium supplemented with appropriate antibiotics for plasmid maintenance and incubated overnight at the appropriate temperature.

### Creation of the *dnaA-PAFP* fusion mutants

The different photoactivatable or photoconvertible proteins were knocked in domain II of DnaA, replacing base pairs 259-312 of the native *dnaA* gene^49,50^. To this end, a previously reported procedure was followed, consisting of a two-plasmid system^48^. Briefly, the wild-type *E. coli* MG1655 cells were made electrocompetent and transformed with the pCas plasmid. Cells were then made electrocompetent again and transformed with one of the four pTarget_dnaA-PAFP plasmids. Correct insertion of the PAFP was confirmed first by PCR using primers BG21364 and BG21365 and then by Sanger sequencing of the locus. The same approach was used for the generation of the *dnaA-PAFP* fusion mutant in the *E. coli* Δ3D background. In this case, the pCas_ampR was used instead of the original pCas plasmid and only the pTarget_dnaA-PAmCherry2.1 plasmid was used to introduce the mutation. After insertion of *PAmCherry2.1* gene in *E. coli* MG1655 and *E. coli* Δ3D, whole genome sequencing was performed to confirm that the modified locus was the only source of DnaA.

In the case of the single deletion mutants, the *E. coli* MG1655 strain carrying the *dnaA-PAmCherry2.1* locus was transformed with plasmid pSIJ8^56^, carrying the λ-red recombineering system. Cells were then made electrocompetent again and transformed with a linear fragment replacing either *DARS1* or *DARS2* with a chloramphenicol resistance cassette or *datA* with a kanamycin resistance cassette. The linear fragments were obtained by amplifying the genomic locus of *E. coli* mutant strains carrying either the Δ*datA::kanR* (primers BG23801 and BG23802), Δ*DARS1::cat* (primers BG23803 and BG23804) or Δ*DARS2::cat* allele^40^ (primers BG23805 and BG23806). Transformant colonies were first selected in the appropriate resistance and the locus of interest was then further confirmed by PCR.

### Turbidostat cultivation of cell cultures

*E. coli* strains were grown overnight in 5 mL of LB medium at 37 °C and 180 rpm. The densely-grown culture was then used to start a turbidostat cultivation using the Chi.Bio platform^57^. Experiments were started according to manufacturer instructions for fluidic lines, electrical connections and user interface. Cultures were started at an OD_600nm_ of 0.05 in M9 medium. Both the M9 supplementation and the temperature were selected according to the specific growth regime to obtain. Default settings were employed for stirring and frequency of optical density measurements. A 650 nm laser diode was used to monitor the optical density of the culture, to minimise accidental photoactivation of PAmCherry2.1. The target optical density was set to 0.4 AU (corresponding to an OD_600nm_ of ∼0.2) and the media reservoir was filled with M9 supplemented as the cultivation chambers. Supplemented M9 medium was pumped in the vessel from the reservoir to maintain a constant optical density. Cells were grown at the target optical density for at least eight generations before the whole culture was harvested for imaging.

In a turbidostat, the dilution rate (*D*) is equal to the growth rate (µ). For each cultivation, the total volume (*V*_SteadyState_) and time (*t*_SteadyState_) spent at the steady-state optical density were used to calculate the flow rate 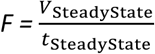. *F* was then used together with the cultivation volume (*V_Culture_* = 20 mL) to calculate dilution rate and thus growth rate, following 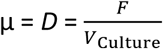.

### Sample preparation and single-particle tracking photoactivatable localisation microscopy

To perform sptPALM of DnaA in live *E. coli,* 10 mL of cells were collected in a 50 mL Falcon tube from turbidostat cultures growing in one of the three different growth regimes (slow, intermediate, fast). Cells were then washed three times in PBS before removing the supernatant and resuspending the pellet in 50 μL of PBS. 1-2 μL of the final cell suspension were immobilised on M9 agarose pads between two heat-treated glass coverslips (#1.5H, 170 μm thickness). The glasses coverslips had been previously heated at 500 °C for 30 min in a muffle furnace to remove organic impurities.

All sptPALM experiments were performed at room temperature using the miCube open microscopy framework^58^. Briefly, the microscope mounted an Omicron laser engine, a Nikon TIRF objective (100x, oil immersion, 1.49 NA, HP-SR) and an Andor Zyla 4.2 PLUS camera running at a 10 Hz for brightfield imaging acquisition and 100 Hz for sptPALM. For each imaging experiment, 300 frames were acquired at 100 ms intervals with brightfield illumination using a commercial LED light (INREDA, IKEA, Sweden). For sptPALM, individual videos of 30,000 frames were acquired at 10 ms intervals. Multiple videos were collected for each field of view, until exhaustion of fluorophores. The Single Molecule Imaging Laser Engine (SMILE) software was used to control the lasers (https://hohlbeinlab.github.io/miCube/LaserTrack_Arduino.html). A 561 nm laser with ∼0.12 W/cm^2^ power output was used for HiLo-to-TIRF illumination with 4 ms stroboscopic illumination in the middle of 10 frames. The 405 nm laser was used to activate the PAmCherry2.1 fluorophores. The 405 nm laser was initially provided with low-power (μW/cm^2^ range) and with a 0.5 ms stroboscopic illumination at the beginning of 10 ms frames^59^. Both the power and the stroboscopic illumination were progressively increased throughout the imaging until exhaustion of fluorophores. Raw data was acquired using Micro-Manager^60^. During acquisition, a 2×2 binning was used, yielding an effective pixel size of 119×119 nm. The excitation field of ∼30×30 µm was restricted to regions of interest of 256×256 pixels or smaller during imaging. The first 500 frames of each video were discarded, to prevent attempted localisation of overlapping fluorophores and to pre-bleach fluorescent contaminants in and around the cells.

### Cells segmentation and cellular area estimation

Fluorophores localisation and cell segmentation were performed as previously described^39^ using ImageJ^61^/Fiji^62^. Briefly, watershed-based segmentation^63^ (http://imagej.net/Interactive_Watershed) was performed to obtain pixel-accurate cell outlines from the brightfield images. From there, a value for the area in pixels of each identified cell was obtained and the camera pixel size (119 nm by 119 nm) was used to generate the corresponding area measurements in µm^2^. The brightfield images, together with their segmented counterparts and list of extracted cell areas is available upon request and will later be available on Zenodo.

### Localisation of fluorophores and tracking of single particles

Single-molecule localisation was performed via the ImageJ/Fiji plugin ThunderSTORM^64^, with added maximum likelihood estimation-based single-molecule localisation algorithms. First, a 50-frame temporal median filter^65^ (https://github.com/HohlbeinLab/FTM2) was applied to the sptPALM movies to correct background intensities^66^. Image filtering was performed through a difference-of-Gaussians filter (Sigma1 = 2 px, Sigma2 = 8 px). The approximate localisation of molecules was determined via a local maximum with peak intensity threshold of *std*(*Wave.F*1) ⋅ 1.2 and 8-neighbourhood connectivity. Sub-pixel localisation was performed through Gaussian-based maximum likelihood estimation, with a fit radius of 4 pixels (Sigma = 1.5 px). A custom-written, MATLAB-based pipeline was used to process and analyse the imaging data. A complete list of localisations for each condition, together with the custom-written MATLAB pipeline is available upon request and will be later available on Zenodo.

Different output files from ThunderSTORM were combined when multiple videos had to be recorded for the same field of view. Localisations were then assigned a cell ID if they fell inside a cell and were discarded if not. Single, valid localisations were linked into tracks according to spatial and temporal distances. The tracking procedure was performed as previously reported^58^ and yielded the number of tracks observed in each single cell and an overall apparent diffusion coefficient distribution. For each track featuring at least 3 localisations, the apparent diffusion coefficient *D*\* was obtained by calculating the mean square displacement between the first *n* steps and taking the average of that, where *n* is the number of localisations minus one. The diffusion coefficients of all tracks were then collected into 85 logarithmic-divided bins from *D\** = 0.04 μm^2^/s to *D\** = 10 μm^2^/s. To investigate the changes in concentration and behaviour of DnaA throughout the cell cycle, the data originated from each strain and growth condition was divided into smaller sets according to cellular area ranges.

### Monte-Carlo diffusion distribution analysis of DnaA

To interpret the diffusional distributions obtained for DnaA-PAmCherry2.1 fusion mutants, a set number of proteins was simulated moving between a bound and a free state in a two linear state model (50,000 for the fit, 250,000 for the visualisation), using Monte-Carlo diffusion distribution analysis (MC-DDA)^39^. The distributions were then fitted with a general Levenberg-Marquardt procedure in MATLAB, yielding kinetic rates with a 95% confidence interval. A single value for localisation uncertainty was used (σ = 0.035 μm, or *D*^∗^_Immobile_ = 0.12 μm^2^/s). DnaA was assigned a diffusion constant *D*^∗^_Free_ = 2.7 μm^2^/s for its free state, based on its hydrodynamic radius and accounting for cytoplasmic retardation due to crowding and viscosity (∼20x)^67^. The value of the DnaA hydrodynamic radius (∼2.48 nm) was obtained using the *in-silico* tool HullRad^68^, providing as input a predicted crystal structure of the DnaA-PAmCherry2.1 fusion mutant, generated through AlphaFold^69^/ColabFold^70^. This diffusion value is similar to previous estimates of diffusion coefficients of proteins moving within the bacterial cytoplasm^71–73^. DnaA was considered as immobile in its DNA-bound state and thus its diffusion coefficient in this state was solely determined by the localisation uncertainty (*D*^∗^ = *D*^∗^ = 0.12 µm^2^/s). Additionally, initial rates *k*_i_ governing the transitions between the free and bound state (*k*_Free→ Bound_, *k*_Bound→Free_) were assigned to the proteins. The proteins were randomly placed inside a cell, simulated as a cylinder (length 2 μm and radius 0.5 μm) with two hemispheres at its poles (radius 0.5 μm). Each protein is randomly put in one of the two different states, based on the probability set by their kinetic rates 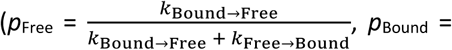 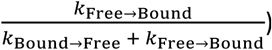. From there, the proteins are given a time before they are changed to a different state, defined as state-change time *t*_Change_ and calculated as 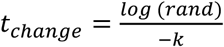, where *rand* − is an evenly distributed random number and *k* is the kinetic rate governing the transition between the current and the next state. The movement of each protein is simulated with over-sampling with regards to the frame-time (*t*_Frame_ = 10 ms, *t*_step_ = 0.1 ms) and their state is recorded every *t*_Frame_ interval. In their free state, each DnaA protein moves for a distance (*s*) equal to a randomly sampled normal distribution, centred around *D*^∗^_Free_ and with 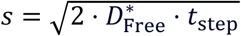. At every step, the *t*_change_ is subtracted with the *t*_step_. If the value becomes ≤ 0, the protein switches its state and new diffusion coefficient and t_change_ are assigned. Every 10 ms, the current location of the proteins is convoluted with a random localisation error, taken from a randomly sampled normal distribution with a localisation uncertainty of σ = 0.035 μm. Each simulated protein had a pre-determined number of localisations, leading to simulated tracks ranging from 1 to 8 steps. The number of tracks of each length follows an exponential decay with a mean track length of 3 steps, following previous experimental observations of PAmCherry2^74^. The bound fraction of DnaA was calculated using the obtained kinetic rates as *p*_Bound_ = 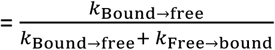. The result was then multiplied by 100 to display it as a percentage.

### Analysis of DnaA behaviour throughout the cell cycle

To study the difference in behaviour of DnaA across the cell cycle, we assigned to each cell a single value of average diffusion coefficient. This value was obtained by summing the diffusion coefficient of tracks a single cell contained and then dividing it by the number of tracks. We then used the area to probe the different phases of the cell cycle, by plotting the obtained average diffusion coefficient over the area of the cell. We then calculated and displayed rolling medians, and 25 and 75% quantiles in R v4.4.1 with the zoo v1.8-12 package^46^, using a rolling window size of ‘k = 75’. Distributions of cell areas were also obtained.

### Flow cytometry estimation of number of origins

To obtain the average number of origins present in an *E. coli* population, both reference cells (wild-type in slow growth regime, 1 to 2 copies) and experimental cells (mutant strains in different growth regimes) samples were prepared^5^. In both cases, turbidostat cultures of *E. coli* were subjected to run-out replication. Rifampicin (150 μg/mL final concentration) and cephalexin (15 μg/mL final concentration) were added, the turbidostat fermentation was stopped and cells were left incubating with constant stirring and temperature for additional 6-6.5 h. Then, 2 mL of the final culture were first washed in the same volume of PBS, subsequently added to 18 mL of ice-cold 70% V/V ethanol and stored at 4 °C for at least 12 h^75^. Reference cells were obtained from turbidostat cultures of wild-type *E. coli* MG1655 in slow growth regime, where cells typically cycle between 1 and 2 copies of the chromosome^4^. Experimental cells were obtained for all strains harbouring the *dnaA-PAmCherry2.1* gene in different growth regimes.

Both reference and experimental cells samples were washed once with PBS and then resuspended in 1 mL of PBS in an Eppendorf tube. The membrane of the reference cells was then stained with MitoTracker^TM^ Deep Red FM (Invitrogen^TM^) to a final concentration of 1 μg/mL and left incubating for 1-1.5 h at room temperature, in the dark, with constant shaking. Reference cells were then washed two times with PBS and finally resuspended in 1 mL of PBS. For absolute DNA content measurement, 25 μL of reference cells were mixed with 75 μL of experimental cells and the cell mixture was stained with 20 μL of 100x Quant-iT^TM^ PicoGreen^TM^ dsDNA reagent (Invitrogen^TM^) in 25% DMSO. The cells were incubated together with the nucleic acid dye for 30-60 min at room temperature in the dark. Finally, 200 μL of PBS was added to the samples.

Analysis of DNA content was performed with an Attune™ NxT Flow Cytometer (Invitrogen^TM^) and the Attune™ proprietary software. For each sample, forward and side scatter measurements were obtained, together with emission filtered with a 530/30 nm transmission filter for DNA content measurement (BL1-H channel, voltage = 260) and emission filtered with a 695/40 nm transmission filter for MitoTracker^TM^ measurement (YL3-H channel, voltage = 450). Reference cells were separated from experimental cells by gating the far-red emission. Thousands of single cells events were collected for the experimental cells. The DNA content of both the reference cells and the experimental cells populations were represented as a histogram versus fluorescence on the green channel. The intensity of the highest peak for the population of reference cells, representing two copies of the chromosomes after run-out replication, was used to quantify absolute number of origins. The intensity maximum of each peak was identified and the number of cells in a 500 A.U. range centred around the peak was used to calculate the average number of origins, asynchrony index and relative abundance of cells with one origin of each sample.

## Results

### DnaA box motifs are preferentially accumulated near the origin of replication

A genomic configuration in which DnaA boxes are distributed preferentially towards *oriC* allows for a large fraction of sites to be replicated soon after the start of DNA replication and is thus ideal for optimal titration of DnaA. Although the current belief is that the hundreds of binding motifs present on the chromosome of *E. coli* are homogeneously distributed^13,20^, this assumption has not been tested yet to our knowledge. We thus decided to compare the number of DnaA boxes per position on the *E. coli* chromosome with the number of boxes expected when sampling from a uniform distribution (Figure 2A). To this end, we used a list of motifs that include both the consensus sequence of DnaA box (TTWTNCACA)^20^, as well as other sequences reported as functional in *datA, DARS1* and *DARS2*^40^. We identified an extended region of significant overrepresentation of DnaA boxes between centisomes 98 and 5, rejecting a uniform distribution and indicating an accumulation around *oriC*. Furthermore, by evaluating the number of exact matches to permutated DnaA box motifs on the chromosome (Figure 2A), we rejected the hypothesis that gradients in genome composition (e.g., GC-content skew) alone could cause the observed enrichment. The same conclusions were reached when only considering the consensus sequence of DnaA box (Supplemental Figure 2A).

**Figure 2.**
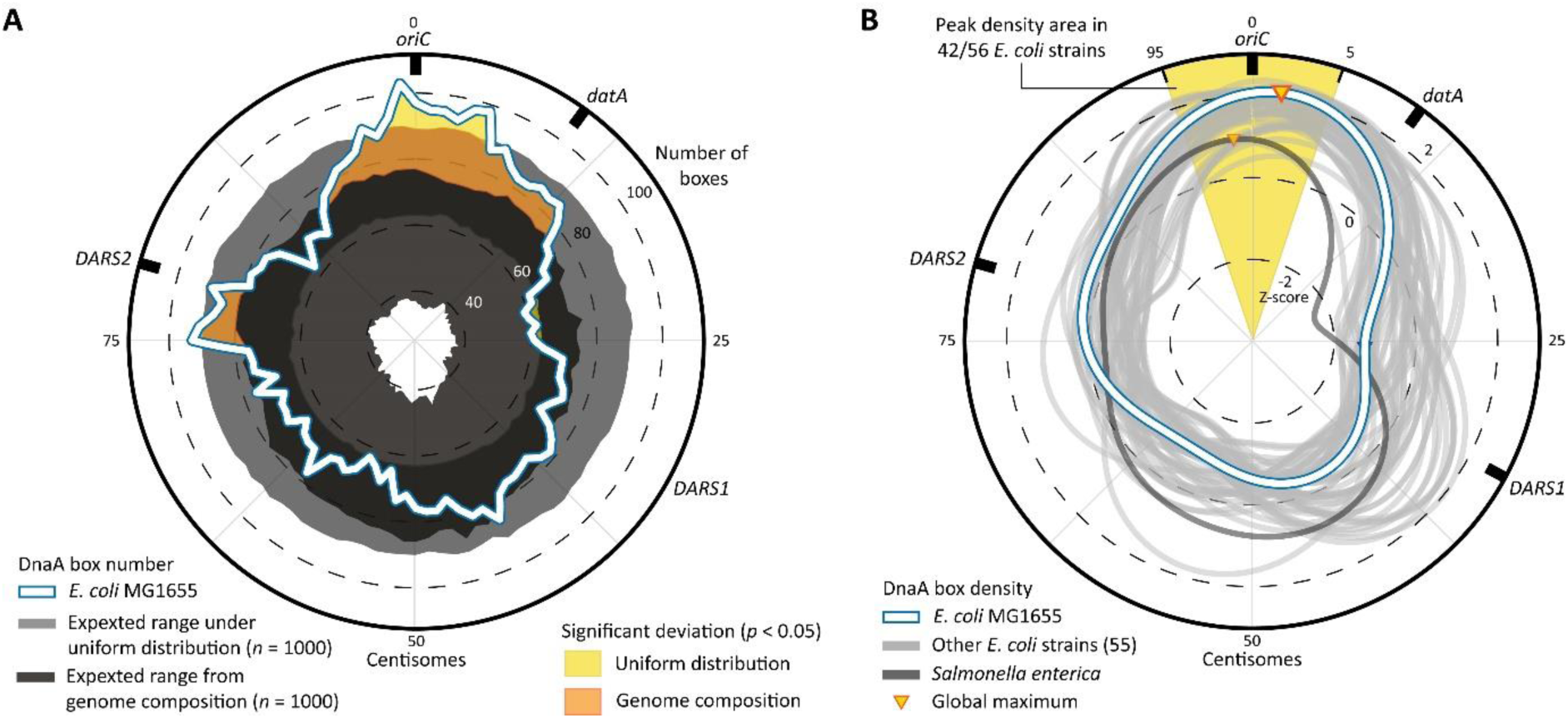
Density peaks of DnaA boxes across the chromosome of *E. coli* strains. **A)** Circular representation of the *E. coli* MG1655 chromosome, depicting DnaA box numbers at distinct positions. The actual numbers of DnaA boxes on the *E. coli* MG1655 chromosome is depicted as the white line and is compared to expected numbers (5 ≤ *p* ≤ 95) based on 1,000 samples from a uniform distribution (light grey bands), and based on sequence matches to 1,000 permutations of DnaA box sequences to account for genome composition biases (dark grey band). The yellow and orange areas indicate regions with significant deviation (pseudo *p*-value < 0.05) from uniform distribution and genome composition, respectively. **B)** Circular representation of DnaA box density across multiple *E. coli* genomes and one singular *Salmonella enterica* chromosomes, expressed as Z-scores. The density of *E. coli* MG1655 is depicted as the white line, while other 55 *E. coli* strains are in grey and *S. enterica*, considered as a distant relative species, is depicted in dark grey. For *E. coli* MG1655 and *S. enterica*, the global maximum is also plotted. See Supplemental Figure 3 for density plots of each individual *E. coli* strain. In all panels, the outer ring represents the relative distance from the origin of replication, expressed in centisomes and with the 0 at *oriC*.

The observed accumulation of DnaA boxes towards *oriC* was not a unique characteristic of *E. coli* MG1655. We found that most *E. coli* chromosomes (n = 42, 73.7% of all analysed strains) have their maximum density of DnaA boxes within 5 centisomes of the *oriC* (Figure 2B). Expanding this analysis, we observed that the global maximum of 86.0% of the genomes peaked within 10 centisomes to *oriC*, with this region having a median DnaA box density two standard deviations above the chromosomal mean (Z-score median = 2.05, interquartile range 1.83-2.15). Interestingly, a similar distribution pattern is also observed in the distant relative *Salmonella enterica* (Figure 2B), suggesting an evolutionary conservation of this genomic arrangement past *E. coli* strains.

Our analysis highlights an accumulation of DnaA boxes near the origin of replication, rejecting previous assumptions on their homogenous distribution^13,20^. The significant overrepresentation of box sequences compared to the genome composition suggests positive selection for DnaA boxes on the chromosome (Supplemental figure 2B) and especially close to the *oriC* (Figure 2A). Furthermore, the preferential accumulation of these motifs on the lagging strand of DNA replication (Supplemental Figure 2C) hints at cooperation between initiator titration and the switch-related mechanism of RIDA^29^, which functions through DNA polymerase III β-clamps^76^ left on the lagging strands of replication^77^. Notably, the abundance of DnaA boxes near *oriC* means that a considerable fraction of DnaA boxes is replicated in the early stages of the DNA replication elongation step. As a result, both the total number of DnaA boxes and their concentration can quickly increase after initiation, a phenomenon helping initiator titration during fast growth^35^.

### Growth-rate-dependent titration of DnaA on the *E. coli* chromosome

After observing that DnaA boxes are preferentially arranged close to *oriC*, we moved to seek experimental confirmation of the ability of the *E. coli* chromosome to titrate DnaA. To this end, we generated several fusions of DnaA with photoactivatable or photoswitchable fluorescent proteins, ultimately selecting PAmCherry2.1^39^ for its low impact on the DNA content and cellular area of *E. coli* (Supplemental Figure 4B-4C). The creation of *E. coli* MG1655 *dnaA-PAmCherry2.1* allowed us to proceed to performing sptPALM (Figure 3A). Building on our previously established framework^39^ (Supplemental Figure 1), we grew the mutant strain at a constant optical density to maintain cells exponentially growing in either a slow (doubling time 187 ± 32 min), intermediate (doubling time 62 ± 9 min) or fast growth regime (doubling time 31 ± 3 min). We then generated the diffusional histograms of the different conditions and fitted them with Monte-Carlo diffusion distribution analysis (MC-DDA) by simulating DnaA moving between a free and a DNA-bound state. In this way, we obtained kinetic rates governing transitions between the two states (Figure 3B). We observed a decreasing trend of the association rate *k*_Free→Bound_ from slow to fast growth regime, whereas the dissociation rate *k*_Bound→Free_ remained constant (Figure 3C, top). Consequently, the overall bound fraction of DnaA decreased from 75 ± 2% in the slow growth regime, to 67± 3% in the intermediate and 55 ± 7% in the fast growth regime (Figure 3C, bottom). The *k*_Bound→Free_ rate extracted from our distribution fittings point at averagely short-lived interactions of DnaA with the chromosome (average *t*_bound_ = 15.7 ± 1.8 ms), consistently with previous reports with lower temporal resolution^78^.

**Figure 3.**
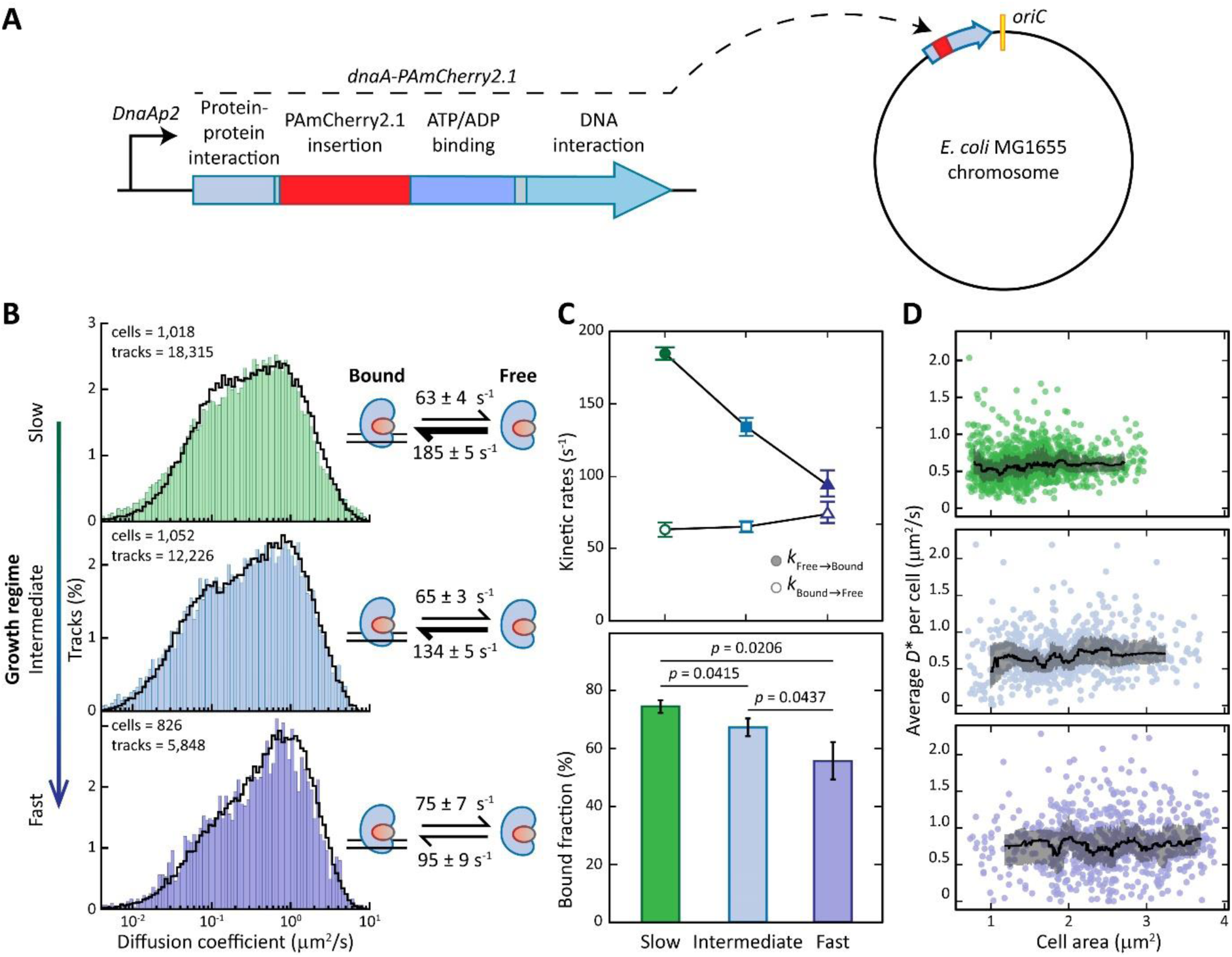
SptPALM of DnaA in live *E. coli* MG1655 cells. **A)** Photoactivatable fluorescent protein PAmCherry2.1 was fused to DnaA in its native chromosomal locus by replacing amino acids 86-113 in Domain II of DnaA. **B)** Distributions of diffusion coefficients of DnaA in live *E. coli* cells grown in either slow, intermediate or fast growth regimes. The number of cells and tracks per histogram is shown in the upper left corner. Histograms are fitted (black line) with a theoretical description of 250,000 particles moving between a free and a bound state. On the right, kinetic rates describing the fitting are provided. **C)** Trends of the kinetic rates *k*_Free→Bound_ and *k*_Bound→Free_ and average bound fraction of DnaA in different growth regimes. **D)** Trends of DnaA mobility throughout the cell cycle. The average diffusion coefficient of each cell is plotted against the area of the cell, with the black line indicating the rolling median of the population. The grey ribbons represent the interquartile range. See Supplemental Figure 4D for the area distribution of the cells.

**Figure 4.**
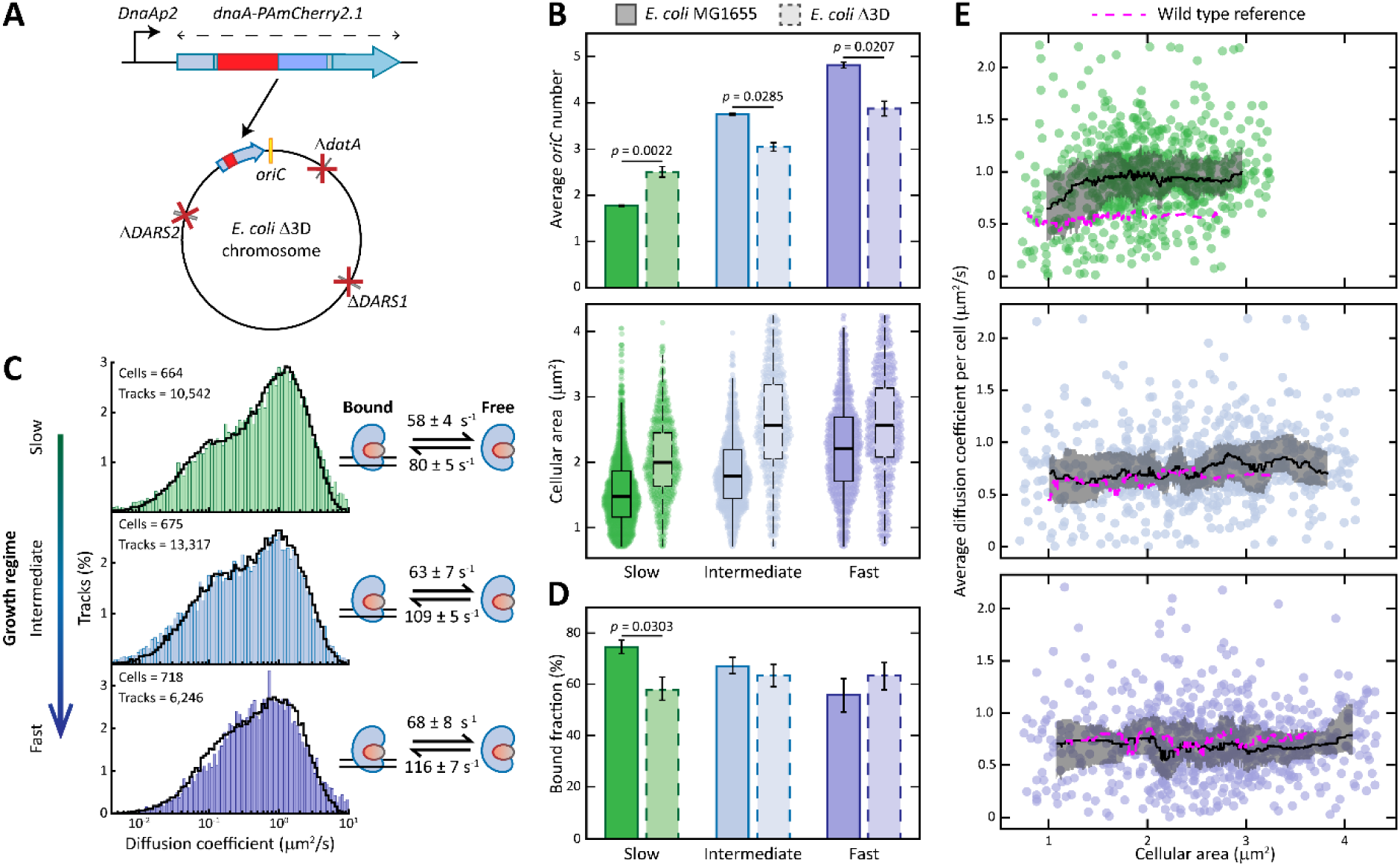
The effect of the Δ*DARS1,* Δ*DARS2,* Δ*datA* triple deletion on the binding of DnaA and the DNA replication of *E. coli.* A) The *dnaA::PAmCherry2.1* mutation was introduced in the chromosomal locus of *dnaA* in the *E. coli* Δ3D strain. **B)** Measurements of DNA content (top) and cellular area (bottom) of either the wild type or the mutant strain for cells grown in slow, intermediate or fast growth regime. See Supplemental Figure 5A for a complete list of replicates of DNA content measurements via flow cytometry. **C)** The diffusion coefficient histograms of DnaA obtained from sptPALM of *E. coli* Δ3D cells growing either in slow, intermediate or fast growth regime. Histograms are fitted (black line) with a theoretical description of 250,000 particles moving between a free and a bound state. The number of cells and tracks per histogram is shown in the upper left corner. On the right, the kinetic rates describing the fitting are provided. **D)** Average bound fraction of DnaA in each growth regime for either the wild-type *E. coli* MG1655 or the triple deletion mutant *E. coli* Δ3D strain. **E)** Trends of DnaA mobility throughout the cell cycle. The average diffusion coefficient of each cell is plotted against the area of the cell, with the black line indicating the rolling median of the population. The grey ribbons represent the interquartile range, whereas the dotted magenta lines represent the wild-type references. See Supplemental Figure 5C for the area distribution of the cells.

We then calculated for each cell an average of the diffusion coefficients of its tracks and compared it with the cellular area. In this way we provided a proxy for the change in mobility of DnaA, indirectly probing its binding state throughout the cell cycle in the given growth condition. This area-dependent analysis further extended our previous observations on the average bound fraction of DnaA to the mobility of this protein in different phases of the cell cycle (Figure 3D). Cells in slow growth regime exhibited a low average diffusion coefficient (∼0.5 μm^2^/s) throughout the whole cell cycle. In intermediate growth regime, cells showed more variation in the average diffusion coefficient median, yet an overall higher DnaA mobility (around 0.7 μm^2^/s). Finally, the overall mobility of DnaA in fast growth regime was constantly the highest throughout the whole cell cycle (∼0.75 μm^2^/s).

Our sptPALM assay revealed that the chromosome can effectively sequester DnaA in a growth rate-dependent fashion, as illustrated by the average DnaA bound fraction and the protein mobility through the cell cycle. This capacity is the highest in slow growth regime and progressively diminishes until a minimum during fast growth.

### *DARS1, DARS2* and *datA* account for a large fraction of the chromosome titration power

Our assay revealed that the chromosome has the capability of sequestering DnaA in live *E. coli* cells during slow growth regime, hinting at initiator titration being a possible mechanism. The genome of this bacterium harbours hundreds of high-affinity DnaA boxes that can recruit both forms of DnaA, peculiarly arranged to have a higher density towards the origin (Figure 2). Whereas the potential function of most of these boxes is still unknown, five of them are at the *oriC*, while 14 more are located among *DARS1, DARS2* and *datA*^40^. These non-coding genomic loci are known to recruit DnaA and play a role in the switch between its active (ATP-bound) and inactive (ADP-bound) forms. To investigate the contribution of these three switch control loci to the state of DnaA, we introduced the *dnaA-PAmCherry2.1* mutation in the triple deletion strain *E. coli* Δ3D (Δ*datA,* Δ*DARS1,* Δ*DARS2*)^40^ (Figure 4A). We then characterised its phenotype at both a population and a single-molecule level.

We first analysed the phenotype of *E. coli* Δ3D, measuring its DNA content (see also Supplemental Figure 5A) and its cellular area in different growth regimes (Figure 4B). The lack of *datA, DARS1* and *DARS2* in fast and intermediate growth regime caused a reduction in DNA content and an increase in cellular area (Figure 4B). The anti-correlation between the average cell size and the average number of origins is consistent with previous reports^18,40^ and with the idea that cells initiate a new round of replication upon reaching a well-defined size^4,5,7^. Interestingly, this anti-correlation is lost when *E. coli* Δ3D is cultured in slow growth regime. The median cellular area increased in this condition (wild-type 1.46 µm^2^, *E. coli* Δ3D 2.19 µm^2^), as observed for intermediate and fast growth, pointing at delayed initiation. However, cells also presented populations with a higher number of origins than the wild-type background. The combination of these two observations suggests that at low growth cells initiated additional, premature rounds of DNA replication in the same cell division cycle. The triple mutant grew similarly to the wild-type strain in slow and fast growth regime and considerably slower in intermediate growth regime (Supplemental Figure 5B). From the number of cells with specific numbers of origins, we also obtained an asynchrony index and observed that this value was higher in *E. coli* Δ3D compared to the wild-type strain in all growth conditions (Supplemental Figure 5D).

We then performed sptPALM on the mutant *E. coli* Δ3D *dnaA-PAmCherry2.1* and calculated diffusion histograms for cells grown in either slow, intermediate or fast growth regime (Figure 4C). Whereas no major changes in the distribution shape or obtained kinetic rates were observed in intermediate and fast growth regime, the absence of *datA, DARS1* and *DARS2* considerably impacted the binding of DnaA during slow growth. As a result, the bound fraction decreased from 75 ± 2% in the wild type to 58 ± 2% in the Δ3D strain in slow growth regime, while remaining similar in intermediate and fast growth regime (Figure 4D). Similarly, following the mobility of DnaA across the cell cycle showed no major changes in cells growing in fast or intermediate growth regime (Figure 4E). Instead, a considerable increase in diffusivity was observed in slow growth compared to the wild-type phenotype, suggesting that *datA*, *DARS1* and *DARS2* account for a considerable fraction of titration power in this regime.

Altogether, the combined deletion of *datA, DARS1* and *DARS2* led to different phenotypes in different growth regimes. The decreased bound fraction and increased mobility of DnaA suggest that the contribution of the three switch control loci on titration is particularly large during slow growth. More interestingly, the higher mobility of DnaA in this condition was also correlated with an increase in the average number of origins of the population. The loss of anti-correlation between the average size and DNA content suggests that at low growth cells are prone to erratic re-initiations of DNA replication during the same cell cycle when titration is altered. Taken together, these results indicate that regulation of the DNA-binding state of DnaA (i.e., bound versus free) is particularly important for the control of replication during slow growth.

### Deletion of single control loci lead to only modest re-initiation of DNA replication

The absence of *datA, DARS1* and *DARS2* during slow growth regime caused a considerable decrease in the DnaA bound fraction. At the same time, this was the only condition in which the triple mutants were not only larger on average, but also featured a higher DNA content. We thus decided to further investigate the behaviour of *E. coli* in slow growth regime. We set out to understand whether the increased availability of free DnaA had an impact in the occurrence of re-initiation events through reduced initiator titration or whether the phenotype could be solely explained through the removal of switch components. To this end, we introduced single deletions of either *datA, DARS1* or *DARS2* in the previously generated *E. coli* MG1655 *dnaA-PAmCherry2.1* strain and compared their phenotype to the wild-type and Δ3D strains.

As before, we performed sptPALM and derived kinetic rates through fitting (Figure 5A). In this way, we estimated the bound fraction of DnaA in the absence of each switch control locus (Figure 5B). The change in bound fraction observed in *E. coli* Δ3D was not a simple sum over the changes in respective single deletion mutants. Only the deletion of *DARS1* caused a reduction in the average DnaA bound fraction (from 75 ± 2% to 69 ± 3%), whereas individually removing *datA* caused an increase to around 80%. Deletion of *DARS2* did not cause a considerable change in the bound fraction of DnaA. By measuring the DNA content of the strains (Figure 5C) we observed an increase in the average number of origins in the absence of *datA* and a decrease of the same value in the absence of *DARS1* when compared to *E. coli* MG1655. These changes indicate that these loci are active during slow growth and are consistent with the idea that removing the inactivator *datA* lowers the initiation volume, while removing the reactivator *DARS1* increases it^35^. The deletion of *DARS2* seemed not to impact the average number of origins of *E. coli*, which remained similar to wild-type levels (1.77 ± 0.004 compared to 1.78 ± 0.016). However, this phenotype was mostly due to the appearance of cell populations with either three or four origins (Supplemental Figure 6A). Instead, the relative abundance of cells with one origin in the *DARS2* deletion mutant was higher than the wild-type reference value (Supplemental Figure 6B), pointing at delayed initiation in agreement with the reactivating function of this locus. *DARS2* was hypothesised to be inactive in slow growth regime^35^ due to the lack of the Fis protein^79–81^. We here report minor activity, perhaps due to the conserved *DARS1* core sequence^40^. The previously described, yet still not fully characterised role of *DARS2* in premature re-initiations^31,81^ can then explain the appearance of cells with three or four origins upon deletion of this locus. Additionally, the average number of origins of the Δ*datA* population was strikingly lower than the one of *E. coli* Δ3D (2.07 versus 2.51), despite a major DnaA re-activator (*DARS1*) still being present. Together with the near-wild-type value of asynchrony index (Supplemental Figure 6C), this result pointed at a low frequency of re-initiation events in in the Δ*datA* strain. Finally, the loss of either *DARS1* or *DARS2* led to cells with a higher area, whereas the deletion of *datA* had the opposite effect (Figure 5C).

**Figure 5.**
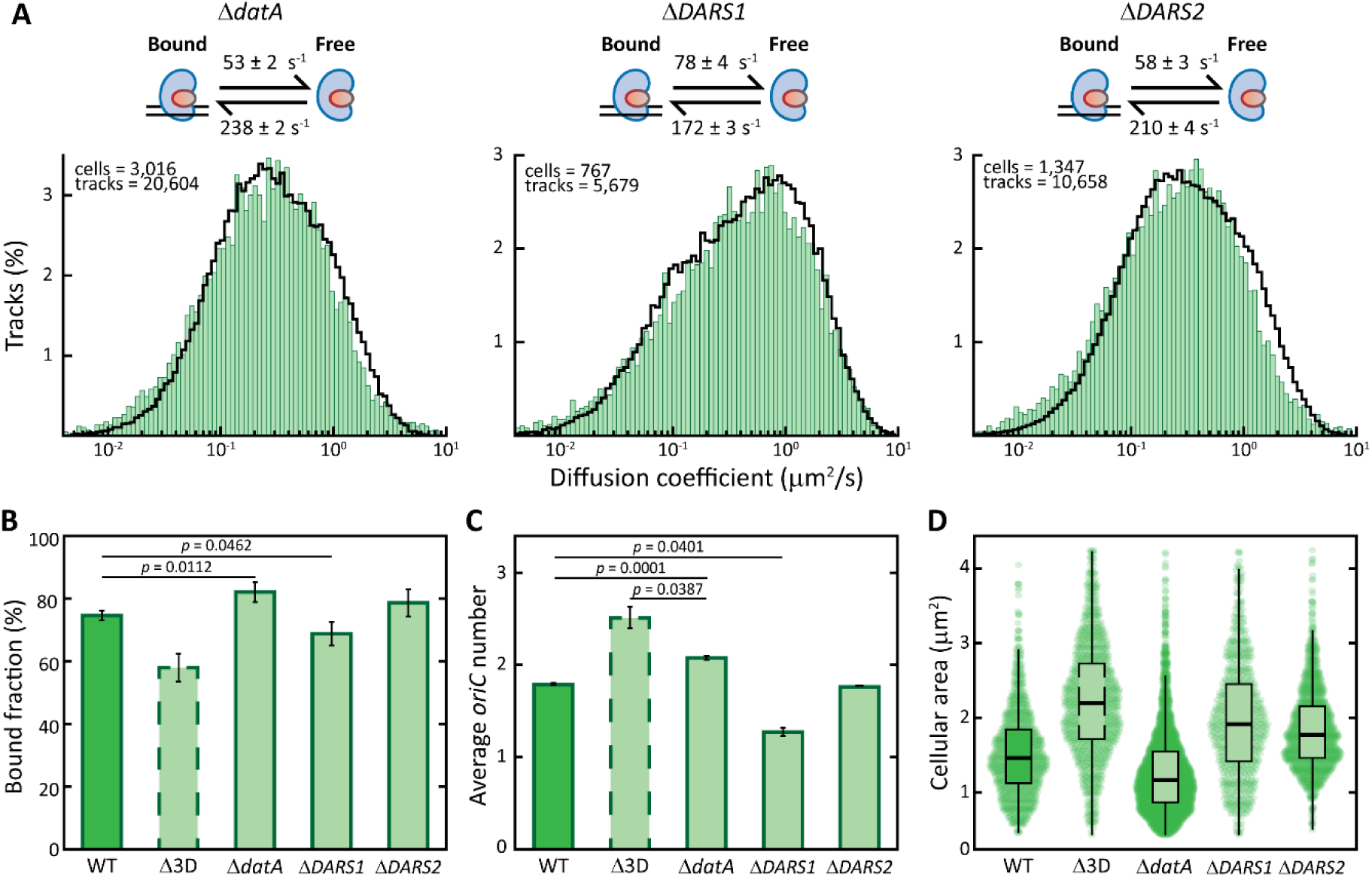
Population- and single-molecule-level phenotype characterization of single *datA, DARS1* or *DARS2* deletions in slow growth regime. A) Diffusion coefficient histograms of DnaA obtained from sptPALM with cells growing in slow growth regime in the absence of either *datA, DARS1* or *DARS2*. Histograms are fitted (black line) with a theoretical description of 250,000 particles moving between a free and a bound state. The number of cells and tracks per histogram is shown in the upper left corner. On top, the kinetic rates describing the fitting are provided for each growth regime. **B)** Average bound fraction of DnaA, as estimated from the kinetic rates, for wild-type, triple deletion and the three single deletion *E. coli* mutants. **C-D)** Comparison of the average number of origins of replication (**C**) and the cellular area of the population (**D**). See Supplemental Figure 6A for a complete list of replicates of DNA content measurements via flow cytometry.

The observed changes in the DNA-bound fraction of DnaA in each of the single deletion mutants are likely a combination of two factors. First, variations in DNA content change the average number of DnaA boxes in the cell compared to the wild-type background. Second, the changes in the DnaA-ATP/DnaA-ADP ratio caused by the deletion of the switch control loci likely affect the binding of DnaA to low-affinity binding motifs on the chromosome, which predominantly occurs for DnaA in its active form^21,22,30^. Overall, deletions of single control loci behaved mostly as predicted by a switch-only model, exhibiting the characteristic anti-correlations between average cell size and average DNA content. These observations are consistent with a model in which replication is initiated once per cell cycle at a well-defined volume, whose threshold value is set by the balance of activation and deactivation^34,35^. The erratic re-initiating phenotype of *E. coli* Δ3D highlighted by our analysis cannot be explained solely by perturbation of factors involved in the switch mechanism. Rather, it revealed a complex behaviour that needs to be explained through a combination of DnaA interconversion and its titration on the *E. coli* chromosome.

## Discussion

In this work we present a systematic characterisation of DnaA behaviour in live *E. coli* cells based on direct visualisation of single DnaA molecules. We confirm previous reports of average transient interactions of DnaA with DNA^78^ with an improved time resolution (10 ms against the previous 41 ms), while further improving on the range of tested growth conditions and genetic backgrounds and implementing an area-dependent analysis. We provide a genomic basis and the first experimental evidence of the titration capacity of the *E. coli* chromosome more than 40 years after it was first proposed^24^. Finally, we show that altering the titration power of the chromosome can impact the control of DNA replication during slow growth.

While assessing the state of DnaA in the wild-type *E. coli* MG1655 (Figure 3), we observed that the association rate *k*_Free→Bound_ constantly decreased from slow to fast growth regime, while the dissociation rate *k*_Bound→Free_ remained roughly constant. As a result, the bound fraction of DnaA was the highest during the slow growth regime and reached a minimum in the fast growth regime (Figure 3C). This phenomenon is the result of two previously described *E. coli* characteristics: (i) balanced biosynthesis^17,18^, leading to constant intracellular DnaA concentration, and (ii) the constant speed of DNA replication elongation^35^. Maintaining constant DnaA concentrations requires increasing the expression of the *dnaA* gene^11,17^ in conditions in which the volume of the *E. coli* cell increases faster, i.e. during higher growth rates^4^. On the other hand, the rate at which new DnaA boxes are replicated per origin is fairly independent of the growth rate, due to the nearly constant speed of DNA replication^5^. Therefore, in increasingly faster growth conditions, the rate of new DnaA synthesis progressively exceeds the rate by which DnaA boxes accumulate^35^. Taking these predictions into consideration, our observed decreasing trend for the association rate indicates an excess of DnaA proteins over the number of available titration sites at the steady-state. As a result, the chromosome ability to maintain a low concentration of free DnaA is impacted at higher growth rates. Still, even at the fastest tested growth conditions (∼30 min of doubling time) the bound fraction accounted for ∼56% of visible DnaA proteins. Pairs of closely spaced boxes have recently been reported to mediate stable binding of DnaA^82^. Therefore, the conserved accumulation of sparse, high-affinity DnaA boxes towards *oriC* on *E. coli* chromosome (Figure 2B) is responsible for the observed bound fractions and supports the existence of a titration-based form of control. With this specific configuration, the number of DnaA boxes can increase quickly enough after DNA replication initiation to always allow more than half of DnaA proteins to be bound. Moreover, the trends of association rates and DnaA bound fractions in different growth regimes are in accordance with previous modelling efforts^35^ postulating the existence of initiator titration as a form of control during slow growth.

The analysis of the triple deletion mutant strain (Figure 4) yielded a complex set of observations that need to be reconciled with existing phenomenological laws. Wild-type *E. coli* initiates one replication round per cell division, an event triggered upon reaching a well-defined initiation volume^4,5^. Genetic modifications that alter the initiation volume but not the growth rate lead to an anti-correlation between the average cell volume and the number of origins^5,7,18,32,40^. We observed such anti-correlation in our *E. coli* mutants lacking a single switch control locus (Δ*datA,* Δ*DARS1* or Δ*DARS2*) during slow growth (Figure 5). The same was true in the *E. coli* triple mutant (Δ3D) only in two of the tested conditions (Figure 4B). The removal of all switch control loci was previously reported to lead to a deficit of reactivating power, due to the leftover presence of the inactivator RIDA^18,40^. Accordingly, *E. coli* Δ3D showed a classical delayed initiation phenotype in intermediate and fast growth regimes, with cells having a lower DNA content and higher average size (Figure 4B). On the other hand, during slow growth, both the average size and number of origins rose over wild-type levels. This phenotype can be interpreted as a mixture of delayed initiation and subsequent premature re-initiation events during the same cell cycle. As in higher growth rates, the lack of enough reactivating power to counteract RIDA^18,40^ led to delayed replication initiation and higher average size. Yet, at low growth rates *E. coli* Δ3D is not capable of lowering its initiation potential enough. Multiple re-initiation events thus occur after the origin of replication is again available for binding^83^, as indicated by the appearance of large populations with three and four origins, the drastic reduction in number of cells with only one origin (Supplemental Figure 5A) and the high asynchrony index (Supplemental Figure 5D). If perturbations to the switch mechanism were the dominant cause of the high frequency of re-initiation events characterising *E. coli* Δ3D during slow growth, a similar phenotype would have also been observed at higher growth rates. The cause of the change in replication behaviour at low growth rates thus likely lies in the alteration of another type of regulation.

Previous modelling efforts reported that an initiator titration mechanism that maintains the free DnaA concentration low throughout the whole cell cycle can enhance the synchrony of replication firing and prevent re-initiation events^33^. In line with this prediction, our area-dependent mobility analysis for the wild-type *E. coli* revealed a constantly low mobility of DnaA from birth to division in the slow growth regime (Figure 3E). Cells then stably cycled between one and two origins, with no observed re-initiation events (Supplemental Figure 5A). In the triple mutant, the deletion of *datA, DARS1* and *DARS2* considerably impacted the binding of DnaA to the chromosome during slow growth, as is evident from the decreased DNA-bound fraction (Figure 4D) and the constantly higher mobility throughout the cell cycle (Figure 4E). Only in this condition cells showed a loss of anti-correlation between size and DNA content, together with a high asynchrony index. Conversely, the mobility of DnaA in *E. coli* Δ3D was unaltered at higher growth rates and cells exhibited a classical phenotype for delayed initiation, with the characteristic anti-correlation between the larger cellular size and the lower number of origins. Therefore, in the slow growth regime, *datA, DARS1* and *DARS2* act as both ‘switchers’, controlling the relative abundance of active and inactive DnaA, and as ‘titrators’, lowering the free DnaA concentration. Removing the three control loci impacted both the interconversion of DnaA and its titration on the chromosome, resulting in the observed premature re-initiations. On the other hand, single deletion of *datA* only altered the interconversion of DnaA but maintained titration intact (Figure 5). The resulting strain showed a classical early initiation phenotype but only modest re-initiations. Thus, our experiments strongly suggest that a reduction in chromosomal titration power does result in firing of subsequent rounds of replication during the same cell cycle.

The double role of *datA, DARS1* and *DARS2* as switchers and titrators prevented us from inferring a role of titration in controlling the first replication event during slow growth. However, our analysis reveals that titration of DnaA through binding to the chromosome does play a role in preventing erratic re-initiations of DNA replication (Figure 6). Titration exerts this effect by ensuring that the free concentration of active DnaA remains low, preventing subsequent, premature binding to the low-affinity boxes on *oriC*. Moreover, titration can enhance the action of RIDA, a DnaA inactivating mechanism. In RIDA, the protein Hda is loaded onto DNA polymerase β-clamps that are left on the lagging strand of DNA replication after the synthesis of Okazaki fragments^77^. From there, Hda inactivates nearby DnaA-ATP by stimulating the hydrolysis of ATP^84^. The striking preference for accumulating DnaA boxes on the lagging strand of each replichore (Supplemental Figure 2C) likely increases interactions with the chromosome in proximity of Hda proteins, ensuring efficient inactivation of DnaA.

**Figure 6.**
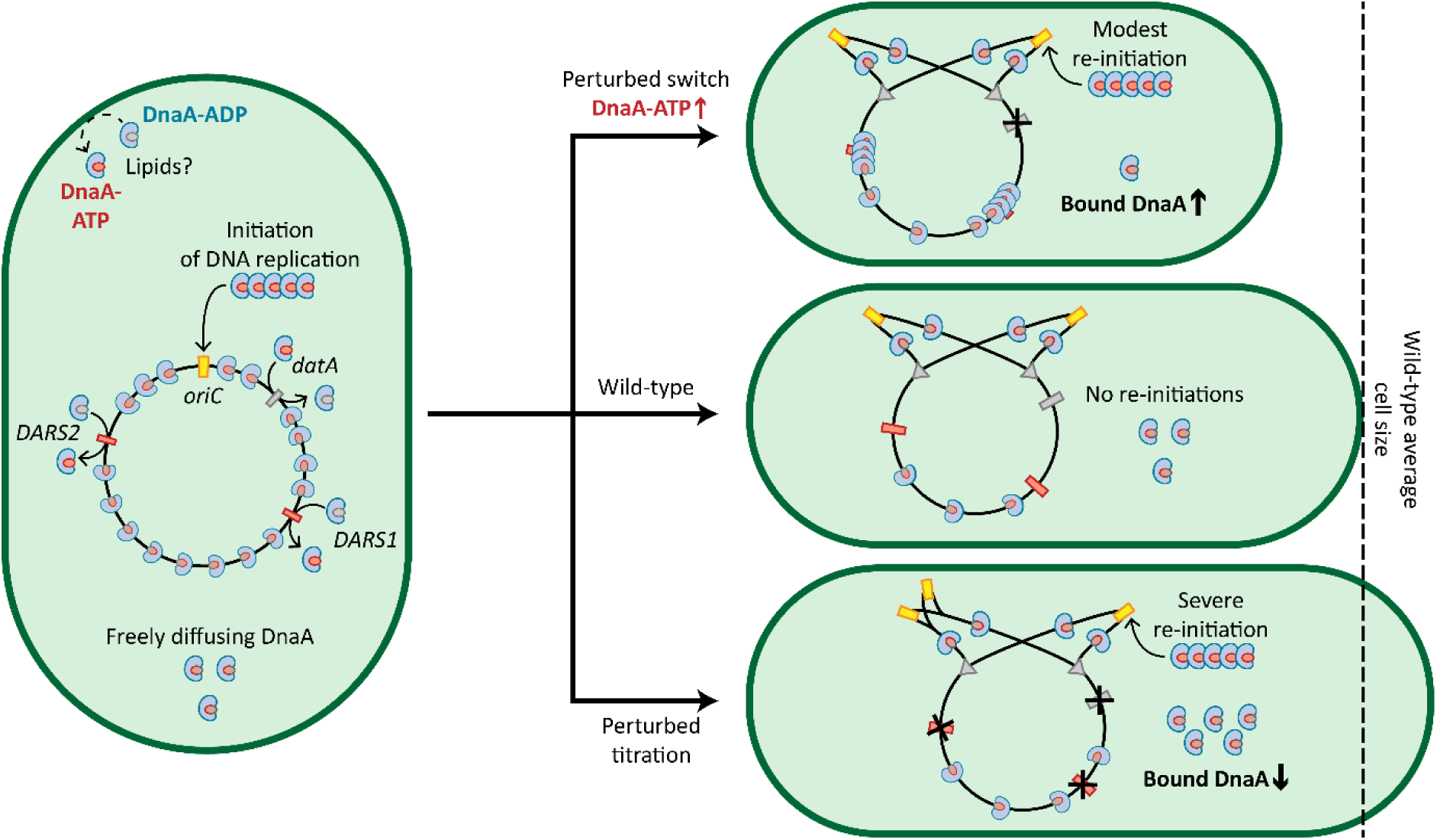
The correlation between initiator titration and re-initiation of DNA replication in *E. coli*. First, DnaA-ATP is accumulated to levels allowing DNA replication initiation (left). The role of titration in the timing of the first replicative event step is still unclear. After DNA replication starts, initiator titration maintains the fraction of active, free DnaA low, preventing re-initiation events (middle). In cases of an altered switch leading to more DnaA-ATP (e.g. *E. coli* Δ*datA*), DnaA titration increases and the frequency of re-initiation events is low (top). On the other hand, cases where titration is hampered (e.g. in *E. coli* Δ3D) lead to a higher amount of free DnaA and severe re-initiations (bottom).

Overall, our findings provide the first experimental support of previous models in which initiator titration stabilises replication cycles^32,35^ and suppresses intrinsic noise in DNA replication initiation. A titration-based control mechanism was hypothesised to be an ancient way of controlling DNA replication in slow-growing ancestors of *E. coli*^32^. Interestingly, the peculiar chromosomal configuration that we found in *E. coli* MG1655, with a high representation of DnaA boxes arranged close to the chromosomal *oriC,* was present in most of the strains of this species, as well as in the relative bacterium *S. enterica*. The conservation of this arrangement perhaps hints at initiator titration being a primordial form of DNA replication control. In the future, specifically altering the titration of DnaA by removing binding motifs outside of *datA*, *DARS1* and *DARS2* could shine further light on the role of this control system also in setting the volume of initiation. In any case, our experimental pipeline will serve as a solid basis for any future research on this elusive form of control, for which direct visualisation of the initiator protein DnaA is posed to be essential.

## Data availability

All data used in this study to generate plots, together with localisation data, the plasmid sequences and the script used in the computational and sptPALM analysis are available upon request by contact with the corresponding authors.

## Acknowledgements

We want to thank Mareike Berger for the inspiring discussions at every stage of the study. We would additionally like to thank prof. Anders Løbner-Olesen (University of Copenhagen) for kindly providing us with the *E. coli* strains carrying deletions for *datA, DARS1* and *DARS2,* alone or in combination. L.O., N.J.C., J.vd.O and P.R.t.W. acknowledge financial support from The Netherlands Organization of Scientific Research (NWO/OCW) Gravitation program Building a Synthetic Cell (BaSyC) (024.003.019). R.H.J.S. is supported by a VIDI grant (VI.Vidi.203.074) from NWO. T.J.G.E. is supported by a European Research Council Consolidator Grant (817834), a VICI grant from the Netherlands Organization of Scientific Research (VI.C.192.016) and a Volkswagen Foundation ‘Life’ grant (96725).

## Notes

### Competing Interest Statement

The authors have declared no competing interest.

## References

1. Willis, L. & Huang, K. C. Sizing up the bacterial cell cycle. Nat. Rev. Microbiol. 2017 1510 15, 606–620 (2017).

2. Helmstetter, C. E. & Cooper, S. DNA synthesis during the division cycle of rapidly growing *Escherichia coli* B/r. J. Mol. Biol. 31, 507–518 (1968).

3. Cooper, S. & Helmstetter, C. E. Chromosome replication and the division cycle of *Escherichia coli* Br. J. Mol. Biol. 31, 519–540 (1968).

4. Wallden, M., Fange, D., Lundius, E. G., Baltekin, Ö. & Elf, J. The Synchronization of Replication and Division Cycles in Individual *E. coli* Cells. Cell 166, 729–739 (2016).

5. Si, F. et al. Invariance of Initiation Mass and Predictability of Cell Size in *Escherichia coli*. Curr. Biol. 27, 1278–1287 (2017).

6. Donachie, W. D. Relationship between Cell Size and Time of Initiation of DNA Replication. Nat. 1968 2195158 219, 1077–1079 (1968).

7. Si, F. et al. Mechanistic Origin of Cell-Size Control and Homeostasis in Bacteria. Curr. Biol. 29, 1760--1770.e7 (2019).

8. Witz, G., Van Nimwegen, E. & Julou, T. Initiation of chromosome replication controls both division and replication cycles in *E. coli* though a double-adder mechanism. Elife 8, e48063 (2019).

9. Le Treut, G., Si, F., Li, D. & Jun, S. Quantitative Examination of Five Stochastic Cell-Cycle and Cell-Size Control Models for *Escherichia coli* and *Bacillus subtilis*. Front. Microbiol. 12, 721899 (2021).

10. Sture Wold, K. S., Steen, H. B., Stokke, T. & Boye, E. The initiation mass for DNA replication in *Escherichia coli* K-12 is dependent on growth rate. EMBO J. 13, 2097–2102 (1994).

11. Zheng, H. et al. General quantitative relations linking cell growth and the cell cycle in *Escherichia coli*. Nat. Microbiol. 2020 58 5, 995–1001 (2020).

12. Katayama, T., Kasho, K. & Kawakami, H. The DnaA cycle in *Escherichia coli*: Activation, function and inactivation of the initiator protein. Frontiers in Microbiology vol. 8 (Frontiers Media S.A., 2017).

13. Hansen, F. G. & Atlung, T. The DnaA tale. Frontiers in Microbiology vol. 9 334195 (Frontiers Media S.A., 2018).

14. Skarstad, K. & Katayama, T. Regulating DNA Replication in Bacteria. Cold Spring Harb. Perspect. Biol. 5, 1–17 (2013).

15. Riber, L., Frimodt-Møller, J., Charbon, G. & Løbner-Olesen, A. Multiple DNA binding proteins contribute to timing of chromosome replication in *E. coli*. Front. Mol. Biosci. 3, 211230 (2016).

16. Dewachter, L., Verstraeten, N., Fauvart, M. & Michiels, J. An integrative view of cell cycle control in *Escherichia coli*. FEMS Microbiol. Rev. 42, 116–136 (2018).

17. Lin, J. & Amir, A. Homeostasis of protein and mRNA concentrations in growing cells. Nat. Commun. 2018 91 9, 1–11 (2018).

18. Boesen, T. O. et al. Dispensability of extrinsic DnaA regulators in *Escherichia coli* cell-cycle control. Proc. Natl. Acad. Sci. 121, (2024).

19. Speck, C., Weigel, C. & Messer, W. ATP– and ADP–DnaA protein, a molecular switch in gene regulation. EMBO J. 18, 6169–6176 (1999).

20. Roth, A. & Messer, W. High-affinity binding sites for the initiator protein DnaA on the chromosome of *Escherichia coli*. Mol. Microbiol. 28, 395–401 (1998).

21. Rozgaja, T. A. et al. Two oppositely oriented arrays of low-affinity recognition sites in *oriC* guide progressive binding of DnaA during *Escherichia coli* pre-RC assembly. Mol. Microbiol. 82, 475–488 (2011).

22. Noguchi, Y., Sakiyama, Y., Kawakami, H. & Katayama, T. The Arg Fingers of Key DnaA Protomers Are Oriented Inward within the Replication Origin oriC and Stimulate DnaA Subcomplexes in the Initiation Complex. J. Biol. Chem. 290, 20295–20312 (2015).

23. Sompayrac, L. et al. Autorepressor Model for Control of DNA Replication. Nat. New Biol. 241, 133–135 (1973).

24. Hansen, F. G., Christensen, B. B. & Atlung, T. The initiator titration model: computer simulation of chromosome and minichromosome control. Res. Microbiol. 142, 161–167 (1991).

25. Katayama, T., Ozaki, S., Keyamura, K. & Fujimitsu, K. Regulation of the replication cycle: Conserved and diverse regulatory systems for DnaA and *oriC*. Nature Reviews Microbiology vol. 8 163–170 (2010).

26. Fujimitsu, K., Senriuchi, T. & Katayama, T. Specific genomic sequences of *E. coli* promote replicational initiation by directly reactivating ADP-DnaA. Genes Dev. 23, 1221–1233 (2009).

27. Sekimizu, K. & Kornberg, A. Cardiolipin activation of DnaA protein, the initiation protein of replication in *Escherichia coli*. J. Biol. Chem. 263, 7131–7135 (1988).

28. Kasho, K. & Katayama, T. DnaA binding locus *datA* promotes DnaA-ATP hydrolysis to enable cell cycle-coordinated replication initiation. Proc. Natl. Acad. Sci. U. S. A. 110, 936–941 (2013).

29. Kato, J. I. & Katayama, T. Hda, a novel DnaA-related protein, regulates the replication cycle in *Escherichia coli*. EMBO J. 20, 4253–4262 (2001).

30. Miyoshi, K., Tatsumoto, Y., Ozaki, S. & Katayama, T. Negative feedback for DARS2 –Fis complex by ATP–DnaA supports the cell cycle-coordinated regulation for chromosome replication. Nucleic Acids Res. 49, 12820–12835 (2021).

31. Frimodt-Møller, J., Charbon, G., Krogfelt, K. A. & Løbner-Olesen, A. DNA Replication Control Is Linked to Genomic Positioning of Control Regions in *Escherichia coli*. PLoS Genet. 12, 1–27 (2016).

32. Knöppel, A., Broström, O., Gras, K., Elf, J. & Fange, D. Regulatory elements coordinating initiation of chromosome replication to the *Escherichia coli* cell cycle. Proc. Natl. Acad. Sci. U. S. A. 120, e2213795120 (2023).

33. Berger, M. & ten Wolde, P. R. Synchronous Replication Initiation of Multiple Origins. PRX Life 1, 013007 (2023).

34. Fu, H., Xiao, F. & Jun, S. Bacterial Replication Initiation as Precision Control by Protein Counting. PRX Life 1, 013011 (2023).

35. Berger, M. & ten Wolde, P. R. Robust replication initiation from coupled homeostatic mechanisms. Nat. Commun. 2022 131 13, 1–13 (2022).

36. Schaper, S. & Messer, W. Interaction of the initiator protein DnaA of *Escherichia coli* with its DNA target. J. Biol. Chem. 270, 17622–17626 (1995).

37. Newman, G. & Crooke, E. DnaA, the initiator of *Escherichia coli* chromosomal replication, is located at the cell membrane. J. Bacteriol. 182, 2604–2610 (2000).

38. Shen, H. et al. Single Particle Tracking: From Theory to Biophysical Applications. Chem. Rev. 117, 7331–7376 (2017).

39. Olivi, L. et al. Live-cell imaging reveals the trade-off between target search flexibility and efficiency for Cas9 and Cas12a. Nucleic Acids Res. 1, 13–14 (2024).

40. Frimodt-Møller, J., Charbon, G., Krogfelt, K. A. & Løbner-Olesen, A. Control regions for chromosome replication are conserved with respect to sequence and location among *Escherichia coli* strains. Front. Microbiol. 6, (2015).

41. Agarwala, R. et al. Database resources of the National Center for Biotechnology Information. Nucleic Acids Res. 46, D8–D13 (2018).

42. R Core Team. R Core Team 2023 R: A language and environment for statistical computing. R foundation for statistical computing. https://www.R-project.org/.RFound.Stat.Comput. 2021 (2023).

43. Blattner, F. R. et al. The Complete Genome Sequence of *Escherichia coli* K-12. Science (80-.). 277, 1453–1462 (1997).

44. Dong, M. J., Luo, H. & Gao, F. DoriC 12.0: an updated database of replication origins in both complete and draft prokaryotic genomes. Nucleic Acids Res. 51, D117–D120 (2023).

45. Agostinelli, C. & Lund, U. R package ‘circular’: Circular Statistics (version 0.5-0). (2024).

46. Zeileis, A. & Grothendieck, G. zoo: S3 Infrastructure for Regular and Irregular Time Series. J. Stat. Softw. 14, 1–27 (2005).

47. Wickham, H. ggplot2: Elegant Graphics for Data Analysis. (Springer International Publishing, 2016). doi:10.1007/978-3-319-24277-4.

48. Jiang, Y. et al. Multigene editing in the *Escherichia coli* genome via the CRISPR-Cas9 system. Appl. Environ. Microbiol. 81, 2506–2514 (2015).

49. Nozaki, S. & Ogawa, T. Determination of the minimum domain II size of *Escherichia coli* DnaA protein essential for cell viability. Microbiology 154, 3379–3384 (2008).

50. Nozaki, S., Niki, H. & Ogawa, T. Replication initiator DnaA of *Escherichia coli* changes its assembly form on the replication origin during the cell cycle. J. Bacteriol. 191, 4807–4814 (2009).

51. Ando, R., Flors, C., Mizuno, H., Hofkens, J. & Miyawaki, A. Highlighted Generation of Fluorescence Signals Using Simultaneous Two-Color Irradiation on Dronpa Mutants. Biophys. J. 92, L97–L99 (2007).

52. De Zitter, E. et al. Mechanistic investigation of mEos4b reveals a strategy to reduce track interruptions in sptPALM. Nat. Methods 2019 168 16, 707–710 (2019).

53. Wang, S., Moffitt, J. R., Dempsey, G. T., Xie, X. S. & Zhuang, X. Characterization and development of photoactivatable fluorescent proteins for single-molecule-based superresolution imaging. Proc. Natl. Acad. Sci. U. S. A. 111, 8452–8457 (2014).

54. Fages-Lartaud, M., Tietze, L., Elie, F., Lale, R. & Hohmann-Marriott, M. F. mCherry contains a fluorescent protein isoform that interferes with its reporter function. Front. Bioeng. Biotechnol. 10, 892138 (2022).

55. Subach, F. V. et al. Photoactivatable mCherry for high-resolution two-color fluorescence microscopy. Nat. Methods 2009 62 6, 153–159 (2009).

56. Jensen, S. I., Lennen, R. M., Herrgård, M. J. & Nielsen, A. T. Seven gene deletions in seven days: Fast generation of *Escherichia coli* strains tolerant to acetate and osmotic stress. Sci. Rep. 5, (2015).

57. Steel, H., Habgood, R., Kelly, C. & Papachristodoulou, A. In situ characterisation and manipulation of biological systems with Chi.Bio. PLOS Biol. 18, e3000794 (2020).

58. Martens, K. J. A. et al. Visualisation of dCas9 target search in vivo using an open-microscopy framework. Nat. Commun. 2019 101 10, 1–11 (2019).

59. Farooq, S. & Hohlbein, J. Camera-based single-molecule FRET detection with improved time resolution. Phys. Chem. Chem. Phys. 17, 27862–27872 (2015).

60. Edelstein, A. D. et al. Advanced methods of microscope control using μManager software. J. Biol. Methods 1, e10 (2014).

61. Schneider, C. A., Rasband, W. S. & Eliceiri, K. W. NIH Image to ImageJ: 25 years of image analysis. Nat. Methods 2012 97 9, 671–675 (2012).

62. Schindelin, J. et al. Fiji: an open-source platform for biological-image analysis. Nat. Methods 2012 97 9, 676–682 (2012).

63. Vincent, L., Vincent, L. & Soille, P. Watersheds in Digital Spaces: An Efficient Algorithm Based on Immersion Simulations. IEEE Trans. Pattern Anal. Mach. Intell. 13, 583–598 (1991).

64. Ovesný, M., Křížek, P., Borkovec, J., Švindrych, Z. & Hagen, G. M. ThunderSTORM: a comprehensive ImageJ plug-in for PALM and STORM data analysis and super-resolution imaging. Bioinformatics 30, 2389–2390 (2014).

65. Jabermoradi, A., Yang, S., Gobes, M. I., Van Duynhoven, J. P. M. & Hohlbein, J. Enabling single-molecule localization microscopy in turbid food emulsions. Philos. Trans. A. Math. Phys. Eng. Sci. 380, (2022).

66. Hoogendoorn, E. et al. The fidelity of stochastic single-molecule super-resolution reconstructions critically depends upon robust background estimation. Sci. Reports 2014 41 4, 1–10 (2014).

67. Trovato, F. & Tozzini, V. Diffusion within the cytoplasm: A mesoscale model of interacting macromolecules. Biophys. J. 107, 2579–2591 (2014).

68. Fleming, P. J. & Fleming, K. G. HullRad: Fast Calculations of Folded and Disordered Protein and Nucleic Acid Hydrodynamic Properties. Biophys. J. 114, 856–869 (2018).

69. Jumper, J. et al. Highly accurate protein structure prediction with AlphaFold. Nat. 2021 5967873 596, 583–589 (2021).

70. Mirdita, M. et al. ColabFold: making protein folding accessible to all. Nat. Methods 2022 196 19, 679–682 (2022).

71. Elf, J., Li, G. W. & Xie, X. S. Probing transcription factor dynamics at the single-molecule level in a living cell. Science 316, 1191–1194 (2007).

72. Persson, F., Lindén, M., Unoson, C. & Elf, J. Extracting intracellular diffusive states and transition rates from single-molecule tracking data. Nat. Methods 2013 103 10, 265–269 (2013).

73. Kapanidis, A. N., Uphoff, S. & Stracy, M. Understanding Protein Mobility in Bacteria by Tracking Single Molecules. J. Mol. Biol. 430, 4443–4455 (2018).

74. Van Beljouw, S. P. B. et al. Evaluating single-particle tracking by photo-activation localization microscopy (sptPALM) in *Lactococcus lactis*. Phys. Biol. 16, 035001 (2019).

75. Ferullo, D. J., Cooper, D. L., Moore, H. R. & Lovett, S. T. Cell cycle synchronization of *Escherichia coli* using the stringent response, with fluorescence labeling assays for DNA content and replication. Methods 48, 8–13 (2009).

76. Kawakami, H., Su’etsugu, M. & Katayama, T. An isolated Hda–clamp complex is functional in the regulatory inactivation of DnaA and DNA replication. J. Struct. Biol. 156, 220–229 (2006).

77. Moolman, M. C. et al. Slow unloading leads to DNA-bound β2-sliding clamp accumulation in live *Escherichia coli* cells. Nat. Commun. 5, 5820 (2014).

78. Schenk, K. et al. Rapid turnover of DnaA at replication origin regions contributes to initiation control of DNA replication. PLOS Genet. 13, e1006561 (2017).

79. Mallik, P., Paul, B. J., Rutherford, S. T., Gourse, R. L. & Osuna, R. DksA is required for growth phase-dependent regulation, growth rate-dependent control, and stringent control of fis expression in *Escherichia coli*. J. Bacteriol. 188, 5775–5782 (2006).

80. Flåtten, I. & Skarstad, K. The Fis Protein Has a Stimulating Role in Initiation of Replication in *Escherichia coli* In Vivo. PLoS One 8, e83562 (2013).

81. Kasho, K., Fujimitsu, K., Matoba, T., Oshima, T. & Katayama, T. Timely binding of IHF and Fis to DARS2 regulates ATP–DnaA production and replication initiation. Nucleic Acids Res. 42, 13134–13149 (2014).

82. Stringer, A. M., Fitzgerald, D. M. & Wade, J. T. Mapping the *Escherichia coli* DnaA-binding landscape reveals a preference for binding pairs of closely spaced DNA sites. Microbiology 170, (2024).

83. Waldminghaus, T. & Skarstad, K. The *Escherichia coli* SeqA protein. Plasmid 61, 141–150 (2009).

84. Riber, L. et al. Hda-mediated inactivation of the DnaA protein and *dnaA* gene autoregulation act in concert to ensure homeostatic maintenance of the *Escherichia coli* chromosome. Genes Dev. 20, 2121–2134 (2006).

