## Supplemental Material for "The *Escherichia coli* replication initiator DnaA is titrated on the chromosome"

**Table S1. List of bacterial strains used in this study.** *DnaA(n1-n2)* stands for the sequence of the *dnaA* gene comprised between base pair n1 and base pair n2, counting from the G at position 1.

| Strains | Characteristics | Source |
|---|---|---|
| *E. coli* MG1655 | K-12; $F^-$ $\lambda^-$ *rph-1* | ATCC |
| *E. coli* Δ3D | MG1655; *strR* Δ*DARS1* Δ*DARS2* Δ*datA* | [1] |
| *E. coli* Δ*datA* | MG1655; *strR* Δ*datA* | [1] |
| *E. coli* Δ*DARS1* | MG1655; *strR* Δ*DARS1* | [1] |
| *E. coli* Δ*DARS2* | MG1655; *strR* Δ*DARS2* | [1] |
| *E. coli* DH5α | MG1655; *endA1 glnV44 thi-1 recA1 relA1 gyrA96 deoR nupG* ϕ80d*lacZ*ΔM15 Δ(*lacZYA-argF)U169 hsdR17* ($r_K^-$ $m_K^+$) | New England Biolabs |
| *E. coli* DH5α x pCas9_ampR | DH5a harbouring the pCas9_ampR plasmid | This study |
| *E. coli* DH5α x pTarget_dnaA-PAFP | DH5a harbouring one of the four plasmids of the pTarget_dnaA-PAFP series | This study |
| *E. coli* MG1655 x pCas9 | MG1655 harbouring the pCas9 plasmid | This study |
| *E. coli dnaA-PAmCherry2.1* | MG1655; *dnaA*Δ*(259-312)::PAmCherry2.1* | This study |
| *E. coli dnaA-dronpa2* | MG1655; *dnaA*Δ*(259-312)::dronpa2* | This study |
| *E. coli dnaA-mEos4b* | MG1655; *dnaA*Δ*(259-312)::mEos4b* | This study |
| *E. coli dnaA-mMaple3* | MG1655; *dnaA*Δ*(259-312)::mMaple3* | This study |
| *E. coli* Δ3D x pCas9_ampR | MG1655 harbouring the pCas9_ampR plasmid; *strR* Δ*DARS1* Δ*DARS2* Δ*datA* | This study |
| *E. coli* Δ3D *dnaA-PAmCherry2.1* | MG1655; *strR* Δ*datA* Δ*DARS1* Δ*DARS2 dnaA*Δ*(259-312)::PAmCherry2.1* | This study |
| *E. coli* Δ*datA dnaA-PAmCherry2.1* | MG1655; Δ*datA dnaA*Δ*(259-312)::PAmCherry2.1* | This study |
| *E. coli* Δ*DARS1 dnaA-PAmCherry2.1* | MG1655; Δ*DARS1 dnaA*Δ*(259-312)::PAmCherry2.1* | This study |
| *E. coli* Δ*DARS2 dnaA-PAmCherry2.1* | MG1655; Δ*DARS2 dnaA*Δ*(259-312)::PAmCherry2.1* | This study |

**Table S2. List of plasmids used in this study.** *DnaA(n1-n2)* stands for the sequence of the *dnaA* gene comprised between base pair n1 and base pair n2, counting from the G at position 1.

| **Plasmid** | **Characteristics** | **Source** |
|---|---|---|
| pCas9 | *repA101*(Ts) *kanR cas9* $P_{araB}$*-Red lacI*$^{q}$ $P_{trc}$-sgRNA-*pMB1* | [2] |
| pSIJ8 | *repA101*(Ts) *ampR* $P_{araB}$*-Red rhaRS* $P_{rhaB}$*-flp* | [3] |
| pCas9_ampR | pCas9; Δ*kanR ampR* | This study |
| pTarget | pMB1; *aadA pJ23119* | [2] |
| pTarget_dnaA-PAFP series | | |
| pTarget_dnaA-PAmCherry2.1 | pTarget; *pJ23119*-sgRNA(*dnaA*) *dnaA(209-258)-PAmCherry2-dnaA(313-362)* | This study |
| pTarget_dnaA-Dronpa2 | pTarget; *pJ23119*-sgRNA(*dnaA*) *dnaA(209-258)-Dronpa2-dnaA(313-362)* | This study |
| pTarget_dnaA-mEos4b | pTarget; *pJ23119*-sgRNA(*dnaA*) *dnaA(209-258)-mEos4b-dnaA(313-362)* | This study |
| pTarget_dnaA-mMaple3 | pTarget; *pJ23119*-sgRNA(*dnaA*) *dnaA(209-258)-mMaple3-dnaA(313-362)* | This study |

**Table S3. List of oligonucleotides and chemically synthesised DNA fragments and their use in the study.**

| Identifier | Sequence (5'-3') | Used for |
|---|---|---|
| BG25452 | ATGTCGGTGATCAAACCAGATATGAAGATCAAACTTCGTATGGAGGGAGCG<br>GTCAATGGGCATCCTTTCGCGATCGAGGGTGTCGGGCTGGGCAAACCCTTCG<br>AAGGGAAGCAAAGTATGGACTTAAAAGTCAAAGAGGGAGGCCCGTTACCCT<br>TTGCGTATGATATTTTGACCACAGTTTTTTGCTACGGGAATCGCGTATTTGCT<br>AAGTACCCGGAAAACATCGTCGATTACTTCAAACAGAGCTTCCCAGAAGGAT<br>ATAGTTGGGAGCGCTCAATGAACTACGAGGACGGTGGCATTTGTAATGCGA<br>CGAACGACATTACATTAGACGGCGACTGCTACATTTATGAGATCCGTTTCGA<br>CGGTGTCAACTTTCCAGCGAATGGACCTGTGATGCAAAAGCGCACTGTAAAA<br>TGGGAACCTTCAACCGAGAAGTTATATGTGCGTGACGGGGTCTTAAAAGGC<br>GATGTAAATACCGCTTTGAGTTTAGAGGGAGGTGGCCATTACCGTTGCGATT<br>TCAAAACGACTTATAAGGCAAAAAAAGTGGTACAATTACCTGATTATCACTTC<br>GTCGATCATCACATCGAGATTAAATCACATGATAAGGACTATTCTAATGTGA<br>ATCTTCACGAACATGCCGAGGCCCATAGTGAGTTACCGCGTCAAGCAAAGTA<br>A | *E. coli* codon-optimised *dronpa2.* |
| BG25454 | ATGGTTAGTGCGATTAAACCAGATATGCGTATCAAGTTGCGTATGGAGGGA<br>AATGTTAATGGACATCACTTCGTCATTGATGGAGACGGAACTGGCAAGCCGT<br>ACGAGGGTAAGCAGACCATGGACCTGGAGGTCAAGGAGGGTGGCCCATTG<br>CCGTTCGCGTTTGATATCCTGACGACAGCTTTCCACTATGGGAATCGTGTCTT<br>CGTAAAATATCCAGATAACATCCAGGACTATTTCAAGCAGTCATTTCCCAAAG<br>GTTACTCTTGGGAACGCAGCTTAACCTTCGAAGACGGGGGGATTTGCAACGC<br>CCGCAACGACATTACTATGGAGGGAGACACGTTTTACAACAAAGTGCGTTTT<br>TATGGAACAAACTTCCCGGCGAACGGGCCTGTTATGCAGAAGAAGACTCTG<br>AAGTGGGAGCCGTCCACGGAAAAGATGTACGTGCGCGATGGGGTCTTAACT<br>GGGGATATTGAGATGGCCTTGCTGCTTGAAGGTAATGCTCACTACCGCTGCG<br>ATTTCCGTACGACCTATAAGGCAAAAGAAAAGGGGGTCAAGTTGCCAGGTG<br>CTCATTTTGTTGATCACGCGATTGAAATTTTGTCGCATGATAAGGATTACAAT<br>AAGGTTAAACTTTACGAGCATGCGGTGGCTCATAGCGGTCTTCCCGACAATG<br>CCCGTCGTTAA | *E. coli* codon-optimised *mEos4b.* |
| BG25455 | ATGGTCTCTAAGGGAGAAGAGACGATCATGTCGGTAATCAAACCGGATATG<br>AAAATCAAGCTTCGCATGGAAGGTAACGTCAACGGTCACGCCTTTGTCATCG<br>AAGGTGAAGGCTCAGGTAAGCCATTTGAAGGTATCCAAACCATCGACTTGG<br>AAGTAAAGGAGGGTGCGCCTTTACCATTCGCATACGACATTCTTACCACCGC<br>TTTCCATTACGGAAACCGCGTGTTCACCAAGTACCCTCGCAAGATCCCTGACT<br>ACTTCAAACAGAGCTTCCCAGAGGGATATTCTTGGGAACGTAGTATGACGTA<br>CGAGGACGGGGGTATCTGCAATGCGACTAATGATATTACAATGGAAGAAGA<br>TTCGTTTATCAATAAAATTCACTTCAAAGGTACGAATTTTCCGCCCAACGGCC<br>CGGTAATGCAGAAACGTACTGTAGGTTGGGAGGTCTCGACTGAAAAGATGT<br>ATGTTCGTGACGGCGTGCTGAAGGGAGATGTTAAAATGAAGCTGCTTCTTAA<br>GGGTGGCTCCCATTATCGCTGTGATTTTCGTACCACATACAAGGTGAAGCAA<br>AAAGCAGTGAAATTGCCCAAAGCACACTTTGTCGACCATCGTATCGAGATCC<br>TTTCTCACGATAAAGATTACAACAAGGTTAAGTTGTATGAACATGCTGTAGCT<br>CGCAATTCCACAGATAGTATGGACGAGCTGTATAAATAA | *E. coli* codon-optimised *mMaple3.* |
| BG21364 | TCAGCCTTAGTCATTATCGAC | Amplified the *dnaA* gene in *E. coli* chromosome |

| | | |
|---|---|---|
| BG21365 | GGTTTACGATGACAATGTTCTG | Amplified *dnaA* gene in all *E. coli* strains |
| BG23801 | CGTATAATGCGCCTCCCG | Amplified Δ*datA::kan* in *E. coli* Δ*datA* |
| BG23802 | CCGAGCCCAAACTGCAAG | Amplified Δ*datA::kan* in *E. coli* Δ*datA* |
| BG23803 | CAGAAAATGCGGCAACCGG | Amplified Δ*DARS1::cat* in *E. coli* Δ*DARS1* |
| BG23804 | AGTTGGGCGGGCAGGTATG | Amplified Δ*DARS1::cat* in *E. coli* Δ*DARS1* |
| BG23805 | GTAAACCACTCTCTGCAGGG | Amplified Δ*DARS2::cat* in *E. coli* Δ*DARS2* |
| BG23806 | GTTGGGACATGTCATGATACC | Amplified Δ*DARS2::cat* in *E. coli* Δ*DARS2* |

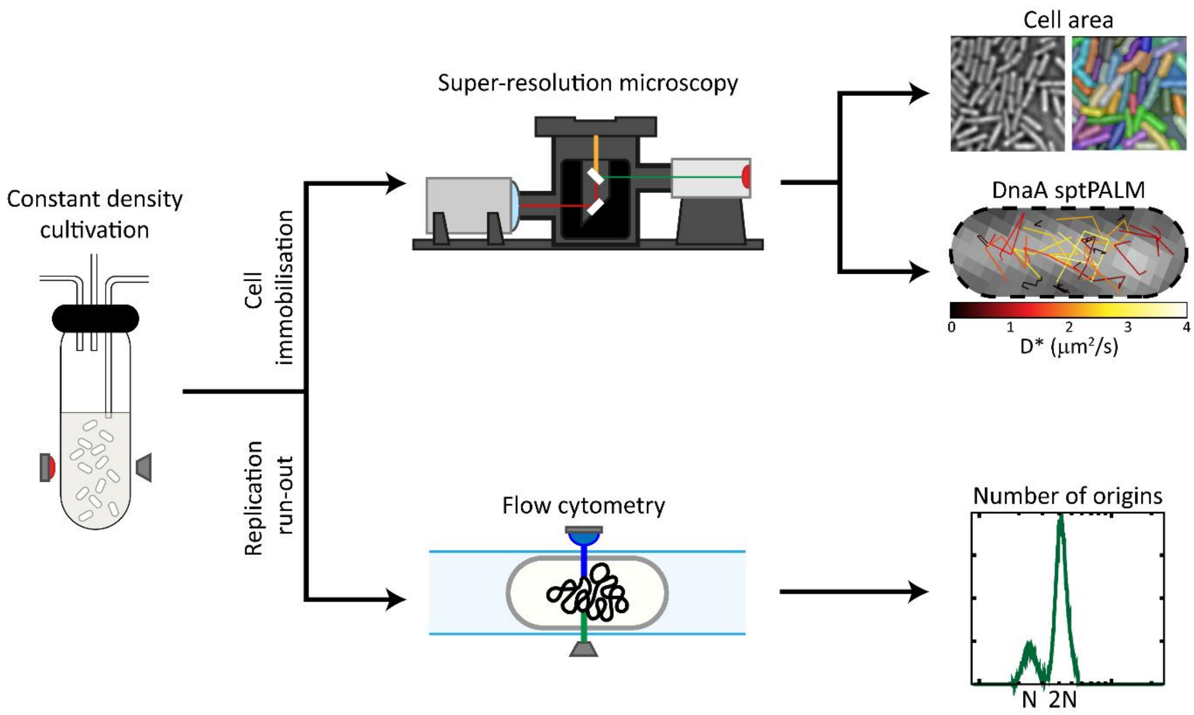


**Figure S1. Experimental pipeline for the single-molecule characterisation of DnaA titration, related to Figure 1.** After an initial growth phase in turbidostat with three different type of media, leading to either slow, intermediate or fast growth regime, cells are processed accordingly to the intended measurements. A first sample (10 mL) is collected, washed in PBS and immobilised on agarose slabs for follow-up microscopy. From the sptPALM imaging, cell area measurements and diffusion coefficients of DnaA tracks were collected. In parallel, replication run-out was performed on the remainder of the cellular culture, by providing rifampicin (150 μg/mL final concentration) and cephalexin (15 μg/mL final concentration) and leave incubating for additional 6 h 30 min at cultivation temperature. A sample of synchronised cells (3 mL) is then collected, fixated and permeabilised in 70% ethanol and then supplemented with PicoGreen before being subjected to flow cytometry for estimating the number of origins of the population.

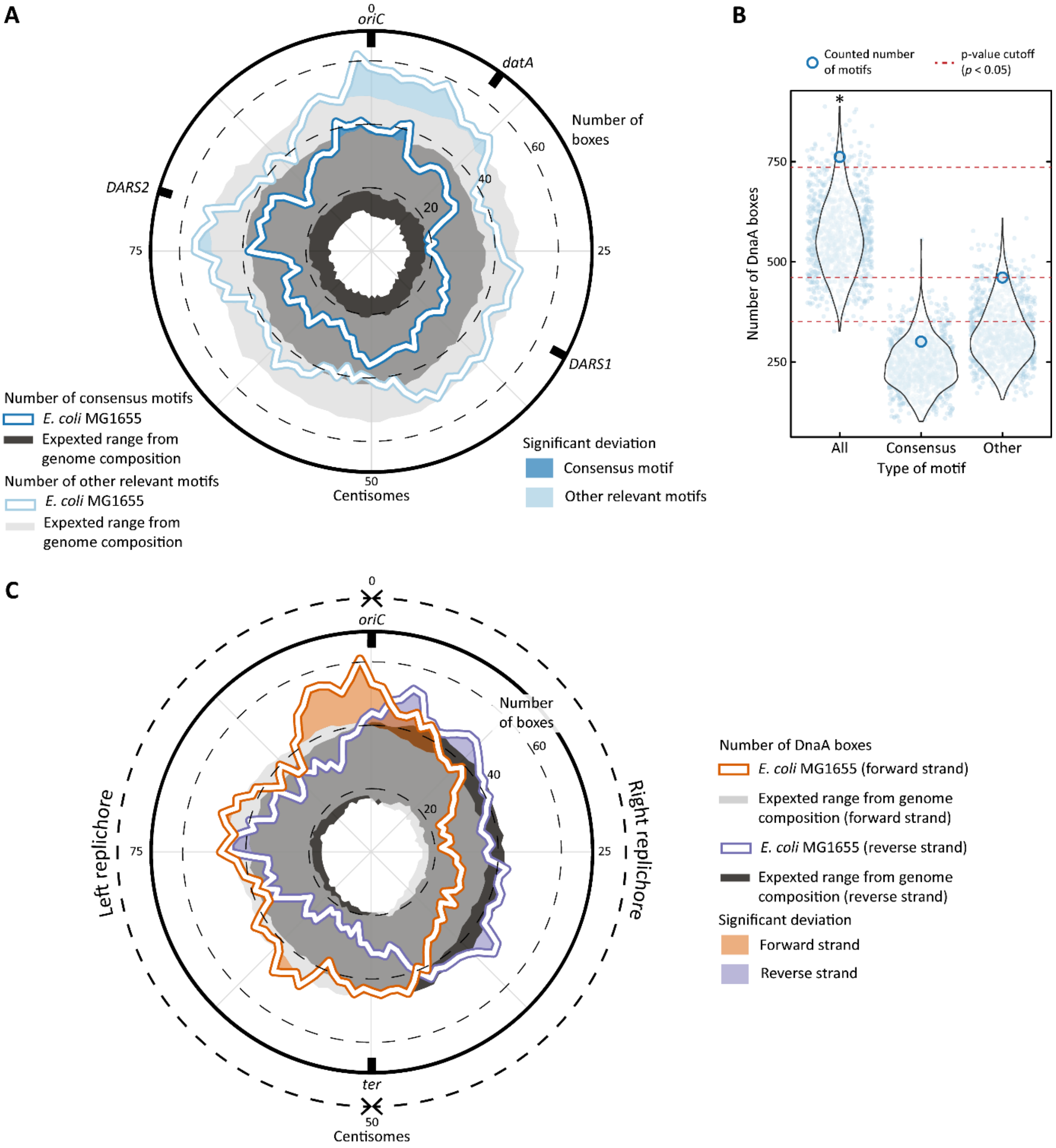


**Figure S2. Computational identification of enrichment of DnaA boxes on the chromosome of *E. coli*, related to Figure 2. A)** Circular representation of the *E. coli* chromosome, with distance units in centisomes. Both the counted consensus DnaA boxes (TTWTNCACA, dark blue) and other sequences (HHMTHCWVH, light blue) are preferentially accumulated towards *oriC,* as observed when compared to the expected range from genome composition. **B)** The total amount of counted DnaA boxes (blue empty dot) is significantly higher than what expected from genome composition (red line indicates threshold of $p < 0.05$). Neither the consensus DnaA box motif nor the other type of motifs (blue empty dots) are individually enriched following the same analysis. The range expected from genome composition was obtained by permutating sequences 1,000 times and counting the instances of each permutated sequence (blue filled dot). **C)** Circular representation of *E. coli* chromosome, with distance units in centisomes. The orange line indicates the boxes counted on the forward strand of *E. coli* MG1655 genome, with coloured areas indicating areas where the count is higher than the expected range from genome composition. Similarly, the purple line and purple area indicate the boxes counted on the reverse strand and enriched areas, respectively. The two replichores are indicated outside: in the right replichore, the reverse strand is the lagging strand during DNA replication, whereas the opposite is true for the left replichore (forward strand is the lagging strand).

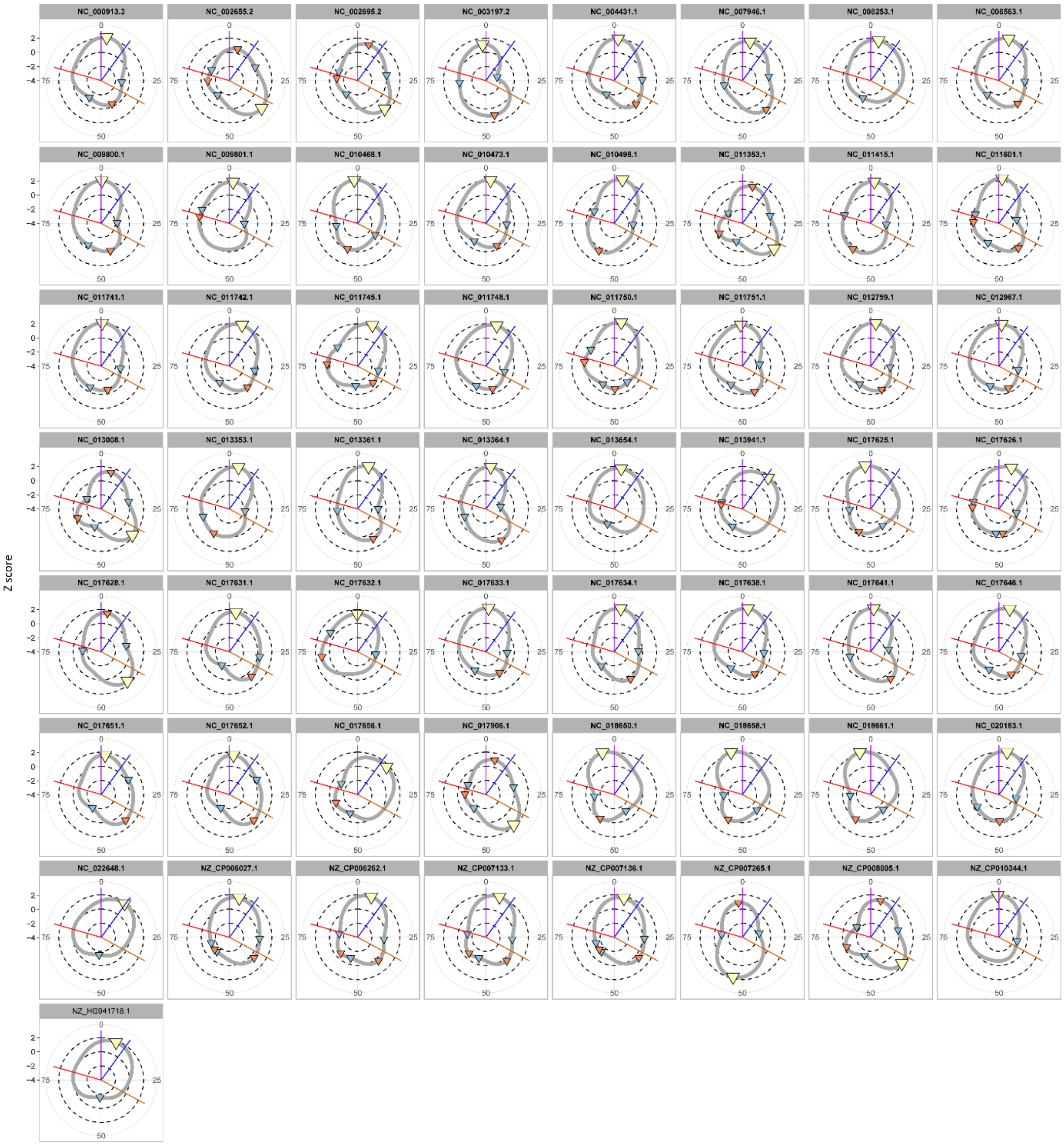


**Figure S3. Individual density plot of each *E. coli* strain with an available sequenced genome, related to Figure 2B.** The accession number of each genome is on top of the related graph. Yellow triangles indicate the global maximum, red triangles indicate local maxima and blue triangles indicate local minima. The purple line indicates *oriC*, the blue line indicates *datA,* the orange line indicate *DARS1* and the red line *DARS2.*

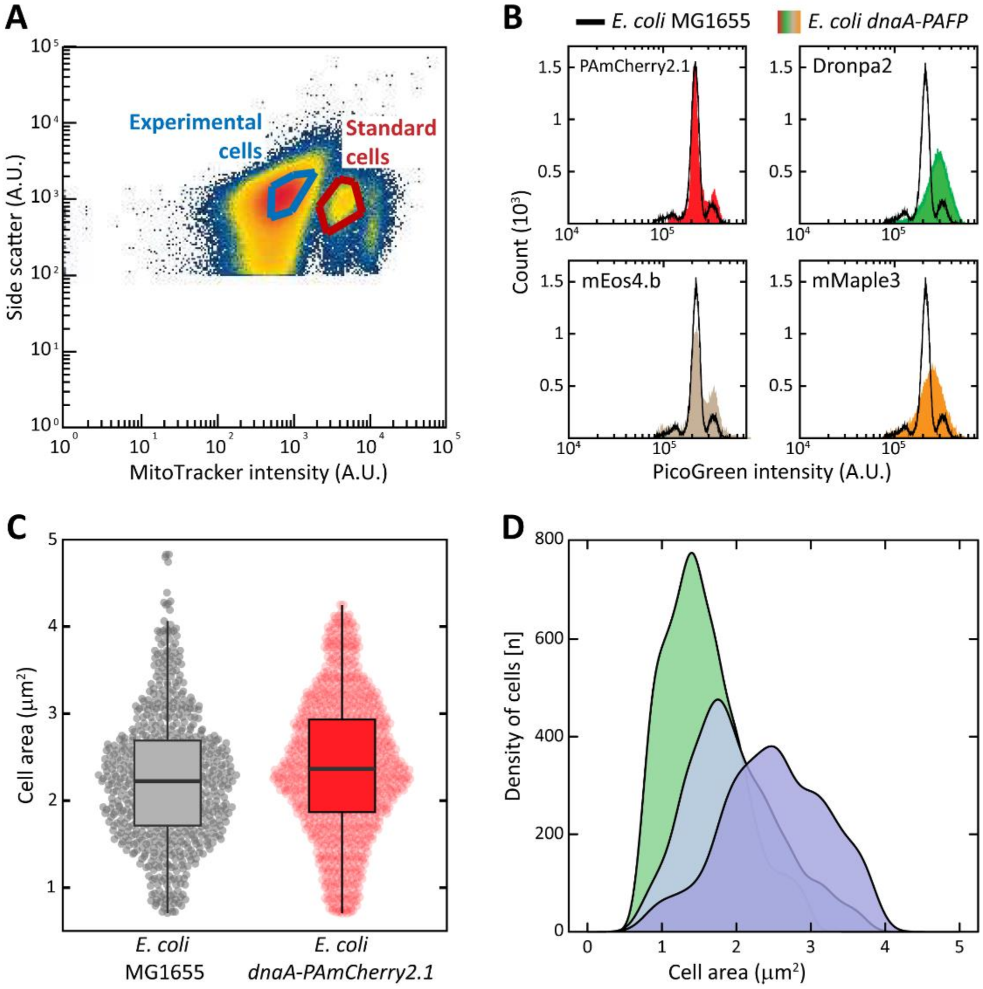


**Figure S4. Characterisation of the mutant *E. coli* MG1655 *dnaA-PAmCherry2.1*, related to Figure 3. A)** Separation of standard and experimental cell by gating for MitoTracker intensity. Cells with a higher far-red emission, derived from the MitoTracker™ dye, were gated as standard cells, whereas the other population was gated as experimental cells. **B)** DNA content of *E. coli* strains carrying fusions of DnaA with either PAmCherry2.1 (red area), Dronpa2 (green area), mEos4.b (beige area) or mMaple3 (orange area) when compared to the wild-type reference (black line). All strains were grown in LB prior to rifampicin run-out and flow cytometry. PAmCherry2.1 impacted DNA replication the least among the tested fluorescent proteins. **C)** Area measurements of the wild type *E. coli* MG1655 and the mutant *E. coli* MG1655 *dnaA-PAmCherry2.1*. All strains were grown in M9 medium, supplemented with 0.4% glucose and 1x RPMI amino acids. The fusion mutant conserved a wild-type cellular area. **D)** Distribution of cellular area of *E. coli* MG1655 *dnaA-PAmCherry2.1* cells grown in either slow (green), intermediate (blue) or fast growth regime (purple) and used for studying the mobility of DnaA across the cell cycle.

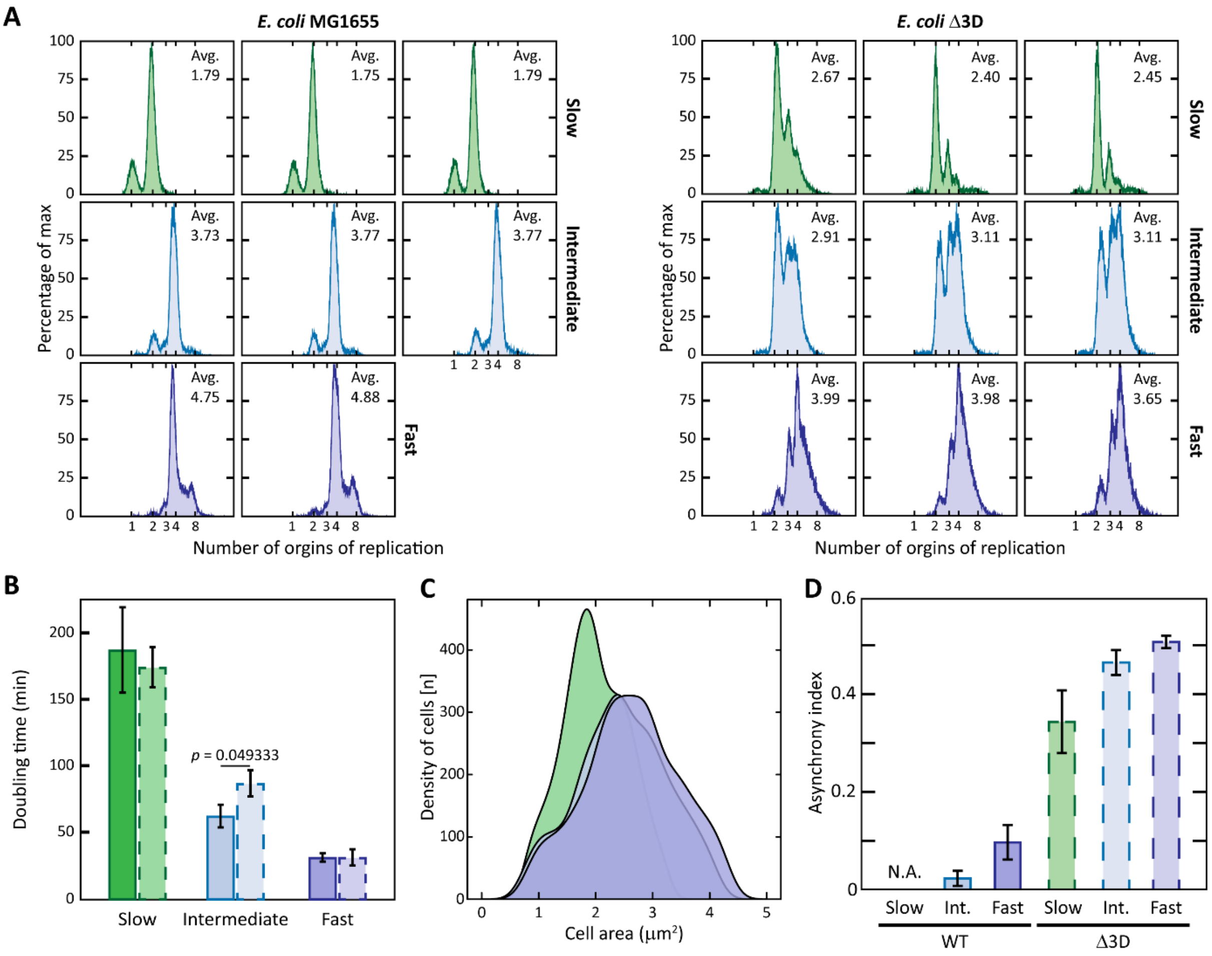


**Figure S5. Cellular parameters used in the study of *E. coli* Δ3D *dnaA-PAmCherry2.1*, related to Figure 4.** In each panel, slow growth regime is indicated in green, intermediate in blue and fast in purple. **A)** Raw data of every replicate of DNA content measurements via flow cytometry of either *E. coli* MG1655 *dnaA-PAmCherry2.1* (left) and *E. coli* Δ3D *dnaA-PAmCherry2.1* (right) in different growth regimes. Average number of origin of replication for each condition is indicated for each replicate. **B)** Growth rate of either *E. coli* MG1655 *dnaA-PAmCherry2.1* (continuous outline) and *E. coli* Δ3D *dnaA-PAmCherry2.1* (dashed outline) in different growth regimes. **C)** Distribution of cellular area of *E. coli* Δ3D *dnaA-PAmCherry2.1* cells in different growth regimes used for studying the mobility of DnaA across the cell cycle. **D)** Asynchrony index of either *E. coli* MG1655, Δ3D, Δ*datA,* Δ*DARS1* or Δ*DARS2.* This value was obtained as Asynchrony index = $\frac{f3+f5+f6+f7}{f2+f4+f8}$, where $f_X$ is the number of cells with X origins of replication[4], as estimated from the replication run-out experiments. Asynchrony index cannot be obtained for *E. coli* MG1655 during slow growth, as cells only cycled between 1 and 2 origins of replication.

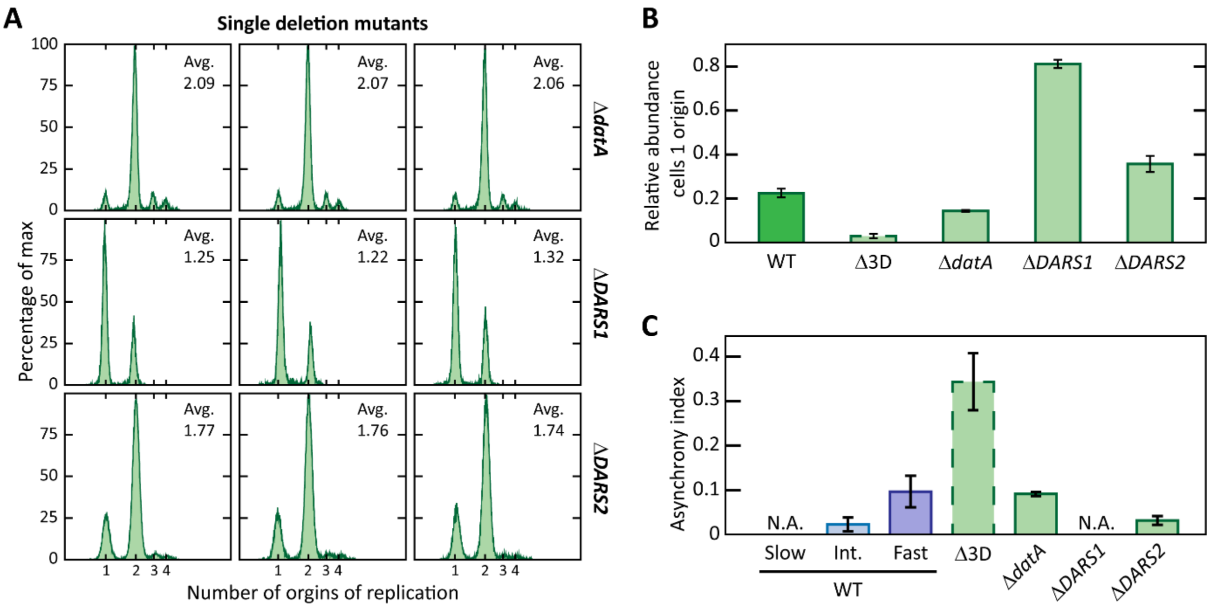


**Figure S6. Cellular parameters used in the study of *E. coli* Δ*datA,* Δ*DARS1* and Δ*DARS2*, carrying the *dnaA-PAmCherry2.1* locus, related to Figure 5. A)** Raw data of every replicate of DNA content measurements via flow cytometry of either *E. coli* Δ*datA dnaA-PAmCherry2.1* (top), *E. coli* Δ*DARS1 dnaA-PAmCherry2.1* (middle) and *E. coli* Δ*DARS2 dnaA-PAmCherry2.1* (bottom) in slow growth regime. Average number of origin of replication for each condition is indicated for each replicate. **B)** The relative abundance of cells with one origin of replication was obtained as $f_1/f_2$, where $f_X$ is the number of cells with X origins of replication, as estimated from the replication run-out experiments. **C)** Asynchrony index of either *E. coli* Δ*datA dnaA-PAmCherry2.1*, *E. coli* Δ*DARS1 dnaA-PAmCherry2.1* and *E. coli* Δ*DARS2 dnaA-PAmCherry2.1*, with *E. coli* MG1655 and *E. coli* Δ3D for comparison. For *E. coli* MG1655, the values obtained for cells grown in intermediate and fast growth regime are provided, as the asynchrony index cannot be calculated for slow growth.

### References

1. Frimodt-Møller, J., Charbon, G., Krogfelt, K. A. & Løbner-Olesen, A. Control regions for chromosome replication are conserved with respect to sequence and location among *Escherichia coli* strains. *Front. Microbiol.* **6**, (2015).

2. Jiang, Y. *et al.* Multigene editing in the *Escherichia coli* genome via the CRISPR-Cas9 system. *Appl. Environ. Microbiol.* **81**, 2506–2514 (2015).

3. Jensen, S. I., Lennen, R. M., Herrgård, M. J. & Nielsen, A. T. Seven gene deletions in seven days: Fast generation of *Escherichia coli* strains tolerant to acetate and osmotic stress. *Sci. Rep.* **5**, (2015).

4. Olsson, J. A., Nordström, K., Hjort, K. & Dasgupta, S. Eclipse–Synchrony Relationship in *Escherichia coli* Strains with Mutations Affecting Sequestration, Initiation of Replication and Superhelicity of the Bacterial Chromosome. *J. Mol. Biol.* **334**, 919–931 (2003).
